# Large seed size accompanying *GmSDP1* suppression is linked to cellular remodeling and phytohormone regulation

**DOI:** 10.64898/2026.11.08.765164

**Authors:** Thiya Mukherjee, Sophia Crowley, Maneesh Lingwan, Kevin L. Chu, Katherine M. Murphy, Katia Gutierrez, Anastasiya Klebanovych, Russell Williams, Anuradha Dhingra, Kirk J. Czymmek, Ben Mansfeld, Timothy P. Durrett, Doug K Allen

**Affiliations:** Donald Danforth Plant Science Center, St. Louis, MO 63132; Department of Natural Sciences, Texas A&M University-San Antonio, San Antonio, TX 78224; Department of Biology, Washington University in Saint Louis, St. Louis, MO 63130; Department of Biochemistry and Molecular Biophysics, Kansas State University, Manhattan, KS 66506; Bayer Crop Science, Chesterfield, MO 63017; Ann &Robert H. Lurie Children’s Hospital of Chicago, Chicago, IL 60611

## Abstract

Soybean (*Glycine max*) seeds contain approximately 20% oil and 40% protein and are important for food, feed, and biofuel. Improving soybean productivity and seed quality requires optimizing carbon partitioning during seed development. Previously, seed-specific suppression of the lipase-encoding gene *Sugar-Dependent 1* (*SDP1*) increased seed oil content by approximately 5% while slightly reducing raffinose family oligosaccharides without compromising protein content. Seed-specific *SDP1* suppression also increased seed size by 18–25% without affecting seed number, resulting in greater seed biomass per plant. To determine the developmental basis of the increased seed size, we investigated the cellular and molecular mechanisms underlying this phenotype. Microscopy and biochemical analyses showed that larger seeds were associated with increased seed cell size and enhanced lignocellulosic cell wall accumulation. Temporal biochemical and transcriptomic analyses further revealed coordinated changes in hormone homeostasis, including altered levels of abscisic acid, 12-oxo-phytodienoic acid, and jasmonoyl-isoleucine, together with differential expression of F-box and RING/U-box family genes. The results indicate that the enhanced seed size resulting from *SDP1* suppression is associated with coordinated changes in cellular development, cell wall accumulation, hormone homeostasis, and regulation of gene expression during seed development. Developmental and metabolic changes associated with the increased seed size phenotype support SDP1 manipulation as a potential strategy for improving soybean seed yield and quality.

## Introduction

Global food demand is increasing with population growth and poses a major challenge to sustainable agriculture and food security (Singh, 2024). The world population is projected to reach 8.5 billion by 2030, resulting in market pressure for increased food, feed, and renewable bioresources (The Sustainable Development Goal report, 2022). Meeting this need will require developing crops with improved productivity and enhanced nutritional value. Because seeds serve as a major source of nutrients and agricultural products, optimizing seed composition in economically important crop species remains a critical objective for crop improvement and agricultural research (Mukherjee et al., 2025).

Soybean (*Glycine max* L. Merr.) is an oilseed crop of high economic importance worldwide. As a legume, soybeans provide a major source of plant-derived protein for human consumption and livestock feed (Allen et al., 2009; Allen and Young, 2013; Yang et al., 2019). As an oilseed, soybean contributes substantially to global plant-based oil production used for food and industrial applications (Allen and Young, 2013). During the 2024/2025 marketing year, soybean represented approximately 62% of global oil seed production, and the United States remained the second-largest producer with a crop value of $43.9 billion (Soystats, 2026). Continued improvements in seed quality and yield are necessary to support agricultural sustainability and economic competitiveness. For the 87 million acres in US soy production, an oil increase by 1% at the expense of undervalued carbohydrate would translate to $1to $2 billion added revenue in agriculture industry (Mukherjee et al., 2025).

Mature soybean seeds typically contain approximately 20% oil and 40% protein, although these values vary depending on genotype and environmental conditions (Litterer et al., 2006; Kambhampati et al., 2021; Aznar-Moreno et al., 2022; Morley et al., 2023). Improving protein and oil simultaneously remains difficult because their levels in seeds are inversely correlated at maturity (Johnson et al., 1955; Hymowitz et al., 1972; Wilcox and Shibles, 2001; Kambhampati et al., 2020). This negative association has long constrained breeding efforts aimed at maximizing seed value. In addition to oil and protein, less valued carbohydrates (30–35%) account for most of the remaining seed biomass (Middelbos and Fahey, 2008). A substantial portion of these includes raffinose family oligosaccharides (RFOs), which are poorly digested by monogastric animals and are considered anti-nutritional components that reduce feed efficiency (Coon et al., 1990; Suarez et al, 1999; Valentine et al., 2017; Aznar-Moreno et al., 2022). Breeding efforts prioritizing yield improvement often negatively affect seed value by decreasing protein and enhancing carbohydrates (Tamagno et al., 2022). These tradeoffs emphasize the importance of understanding carbon allocation among competing storage pathways during seed development. Carbon partitioning and seed metabolism dictates the balance between lipid, protein, and carbohydrate accumulation and ultimately determines seed composition and value (Kambhampati et al., 2026). Developing strategies to redirect carbon flux toward high-value seed reserves will provide opportunities to improve soybean nutritional quality while maintaining productivity.

Seed development in soybean follows an ordered progression through three distinct phases of embryogenesis: early development (morphogenesis), mid development (cell enlargement and seed filling), and late development leading to desiccation and maturation (Thien Nguyen et al., 2016; Mukherjee et al., 2025) Final seed composition is restricted by the resources the seed receives throughout development and the metabolic flux through pathways that convert these resources to seed reserves (Allen and Young, 2013).During the seed-filling phase, maternal carbon inputs (sugars and amino acids, (Allen and Young, 2013; Pegler et al., 2023; Mukherjee et al., 2025)), together with carbon released from starch turnover support the continued synthesis of lipids, proteins, and carbohydrates, including RFOs and cell wall polysaccharides. By late-seed development, the supply of vegetative carbon through the phloem has ceased and developing seeds partially remobilize stored lipids and proteins to sustain carbohydrate or other biosynthetic demands (Chia et al., 2005; Baud et al., 2008; Baud and Lepiniec, 2009; Kambhampati et al., 2021; Mukherjee et al., 2025).

One strategy to improve seed oil accumulation is to minimize lipid degradation during late seed development. Seed-specific suppression of the triacylglycerol (TAG) lipase Sugar-Dependent 1 (SDP1) through RNA interference (RNAi) has been shown to increase total seed oil content by limiting TAG turnover (Kelly et al., 2013; Kim et al., 2014; Kanai et al., 2019; Azeez et al., 2022; Aznar-Moreno et al., 2022). Prior work demonstrated that suppression of *SDP1* in soybean seeds increases fatty acid accumulation at maturity; primarily through reduction of RFO production while maintaining total protein content (Aznar-Moreno et al., 2022). Consistent with earlier observations, enhanced oil accumulation resulting from *SDP1* suppression did not negatively affect seed maturation or germination (Kelly et al., 2013; Kim et al., 2014; Azeez et al., 2022; Aznar-Moreno et al., 2022).

In addition to restricting late-stage seed TAG turnover and enhancing seed oil accumulation, suppression of *SDP1* has consistently been associated with increased seed weight across multiple species. Previous studies reported an increase in seed weight ranging from 13–30% in *Camelina sativa* (Dhankher, 2021), 16% in the *Arabidopsis thaliana sdp1* mutant (Kim et al., 2014), 12–16% in *Physaria fendleri* (Azeez et al., 2022), and 4–14% in *Glycine max* (Kanai et al., 2019). Our prior work demonstrated a 14–23% increase in seed weight in soybean RNAi *SDP1*-suppressed lines, which, together with elevated fatty acid accumulation, resulted in an overall 20% increase in oil yield relative to wild type (Aznar-Moreno et al., 2022). Despite these consistent phenotypic gains, the biochemical and cellular mechanisms underlying enhanced seed weight following *SDP1* suppression remain unknown.

In this study, we investigated the developmental basis of the increased seed weight phenotype resulting from *SDP1* suppression. Increased seed weight became evident during mid seed filling and persisted through maturation. This increase was associated with greater seed size, enhanced cell expansion, and increased cell wall accumulation, without changes in seed number. Developmental transcriptomic and biochemical analyses further implicated changes in phytohormone signaling and the ubiquitin–proteasome system during seed development. Together, these findings provide insight into the developmental and metabolic changes associated with suppression of seed lipid turnover and support *SDP1* as a potential target for increasing seed oil content and yield while maintaining seed quality in soybean and other species.

## Material and Methods

### Plant growth conditions and sample preparation

Soybean seeds (*Glycine max* cv Williams 82) were planted in one-gallon pots with Berger BM7 35% bark soil (10121500; Hummert, Earth City, MO, USA) and grown in a greenhouse supplemented with 14h day: 10h night photoperiod with 25°C/23°C Day/night temperature and 50% relative humidity. Seeds were sampled across development as described before (Aznar-Moreno et al., 2022). Briefly, pods were harvested at three reproductive stages (R5.5, R6, and R7) and at maturity (R8). Seeds were immediately harvested, flash-frozen in liquid nitrogen, and stored at –80 °C until further use. Prior to analysis, seed coats were removed, and cotyledons were ground using an in-house stainless-steel hammer-crushing pestle and mortar. The ground, frozen tissue was aliquoted, with other fractions kept fresh or lyophilized for subsequent analyses.

### Phenotypic and agronomic measurements

Four to six plants for wild-type (WT) and each of the threeT4 homozygous transgenic genotypes; A (*sdp1-1 RNAi A*), B (*sdp1-3, RNAi D*) and C (*sdp1-3, RNAi C*) (Aznar-Moreno et al., 2022) were grown in one-gallon pots, randomized within the greenhouse. Phenotypic traits, including node and pod number were recorded when approximately 95% of the pods had reached maturity, as indicated by dry yellow color and the absence of new pod growth. Seeds were hand-harvested, and a seed count per plant was determined. Average seed weight was determined by weighing 100 seeds individually from each of four to six plants per genotype. Average individual seed area was obtained from ten seeds each from four to six plants per genotype. Red-green-blue (RGB) images were captured using a standard digital camera (Nikon Coolpix L830) connected to Raspberry Pi computer for capturing, labeling, and transferring the images, as previously described (Duenwald et al., 2025). Bright, indirect lighting was used, and a color card was used for color correction and size standardization. Images were processed using open-source software PlantCV (Gehan et al., 2017). First, images were color corrected, and then seeds were masked (segmented) from the background. Individual seeds were measured using the morphology package for seed size and shape **(Supplementary figure S1D)**. To capture an image of all seeds for visual comparison **(Supplementary figure S1C)** the same protocol was used, except a Nikon D7200 was used instead of a Coolpix.

### Microscopy analysis

#### Sample Preparation

Three to five mature soybean seeds were fixed in 4% paraformaldehyde (15710; Electron Microscopy Sciences, Hatfield, PA, USA and Leica Microsystems, Wetzlar, Germany) in 1X PBS buffer (phosphate buffered saline pH 7.4, 137 mM NaCl, 2.7 mM KCl, 10 mM Na_2_HPO_4_, 1.8 mM KH_2_PO_4_), (19342-10; Electron Microscopy Sciences, Hatfield, PA, USA) for 48 hours at 4°C. After fixation, samples were thoroughly rinsed in 1XPBS buffer, briefly dried and embedded in O.C.T. compound cryo-embedding medium (62550-0; Electron Microscopy Sciences, Hatfield, PA, USA). Each sample was sectioned using Leica CryoJane CM1950 cryostat (Leica Biosystems, Buffalo Grove, IL, USA) to expose the tissue structure. This step was done to avoid hand-free razorblade sectioning and ensure the exposed tissue surface was even and flat. Sectioned seeds were incubated in 25 μM Calcofluor White fluorescent dye (29067; Biotium, Fremont, CA, USA) to stain cell walls. After rinsing, samples were mounted in distilled water on the No.1.5H glass-bottom WillCo-dish® (HBST-5040; Willco Wells, Amsterdam, The Netherlands) designed for inverted microscope confocal imaging.

#### Imaging

Each biological replicate was imaged using the same acquisition conditions as described below. 4-10 fields of view (FOV) were captured per replicate using Leica TCS SP8-X laser confocal microscope (Leica Microsystems, Wetzlar, Germany) equipped with 405-nm UV laser, HC PL APO CS 20X/NA 0.7 dry objective, pinhole 1 AU, and PMT detector. Each FOV was acquired as a 12-bit, either 2D 1024×1024 pixels image with the pixel size of 0.568 × 0.568 μm, or in some cases as a 5-μm z-stack with a z-step size of 2 μm and a voxel size of 0.568 × 0.568 × 2 μm. To detect a cell wall stained with the Calcofluor White fluorescent dye (Excitation /Emission 360/430 nm), a UV laser (405 nm) was used with Acousto-Optical Beam Splitter prism set to the emission detection of 415-470 nm.

#### Image processing and data analysis

Cell area and cell count were analyzed using the Fiji open-source software (Schindelin et al., 2012). All images were processed as 2-dimentional (2D) or Maximum Intensity Projections (MIPs) of 1024 × 1024-pixel FOV using the Trainable Weka Segmentation plugin (Arganda-Carreras et al., 2017). This approach enabled segmentation and selection of the inner regions delineated by cell wall staining. A binary mask was then generated, followed by the application of “Fill holes” and “Watershed” filters. Subsequently, the Analyze Particles plugin was used to calculate the total area of each cell in μm^2^ and determine the total number of cells per 1024×1024 pixel FOV. Cells that were incomplete or in contact with the frame borders were excluded from the analysis. Density distribution curves of cell area were plotted using OriginPro software (2024; OriginLab, Northampton, Massachusetts, USA).

### Fresh weight and dry weight measurements of developing seeds

Seeds were harvested from four individual plants per genotype for each of the three *sdp1* suppressed lines and the WT control at three developmental stages: R5.5, R6, and R7. Fresh weight (FW) and dry weight (DW) were determined using an analytical balance (AG204 Delta Range, Mettler Toledo, Columbus, Ohio, USA). For FW measurement, the weight of individual 2 mL microcentrifuge tubes was first recorded. A single seed was then placed in each tube, and the combined weight was measured. Seed FW was calculated by subtracting the pre-recorded empty tube weight from the total weight. Immediately following FW determination, seeds were placed in a drying oven set at 60°C. Dry weight was measured at 3-day intervals over a 20-day period, until no moisture remained as indicated by a stable DW value.

### Quantification of metabolites across development

Total lipids, starch, and oligosaccharides were quantified from three to four biological replicates per reproductive stage. To minimize the effects of canopy position, seeds were collected from multiple positions within the canopy of each plant replicate, pooled, and ground together. For each plant replicate, two to four technical replicates were quantified.

#### Fatty Acid Methyl Ester (FAME) analysis for lipid content

10-15mg of lyophilized tissue was weighed into 8ml glass tubes and analyzed using 1 mg of C17:0 TAG internal standard (T2151; Millipore Sigma, Saint Louis, MO, USA). To each sample, 2ml of freshly prepared 5% (v/v) sulfuric acid: methanol was added. To prevent oxidation of lipids, 25ul of 0.2% butylated hydroxytoluene (101162; MP Biomedicals, Irvine, CA, USA) in methanol was added. Reactions were incubated on a heat block at 110°C for three hours with intermittent vortexing, initially after 20 mins and then after each hour interval. After three hours, tubes were cooled at room temperature, and the reaction was quenched using 3ml of 0.9% sodium chloride. FAMEs were extracted using 1ml hexane and quantified by gas chromatography-flame ionization detection (GC-FID) using a DB23 column (30m,0.25mm i.d.,0.25 um film; J and W Scientific, Folsom, CA, USA). 20:1 split mode was used in the GC. Flame detector temperature was set to 250°C with the oven temperature ramped from 170°C to 250°C at a rate of 10°C /min with a program hold time of 6 min respectively. Quantification was performed using peak area comparison with internal standard.

#### Starch quantification

Starch extraction and quantification was performed using the Megazyme starch assay kit (Megazyme International, Ireland), AOAC Official Method 996.11 (Mccleary et al., 1997; McCleary et al., 2019) with assay volumes adjusted for 96-well plate use as described previously (Kambhampati et al., 2021). Briefly, 15-25mg of lyophilized tissue was washed twice with 80% (v/v) ethanol at 85°C, incubated with DMSO at 110°C for 10 min, digested with α-amylase at 110°C for 12 min with intermittent vortexing, and digested with amyloglucosidase at 50°C for 1 hr. Samples were then centrifuged and the aliquots of the supernatant were incubated with the GOPOD reagent mixture at 50°C for 20 mins before absorbance at 510 nm was measured. Starch concentrations were determined using standard curves of serially diluted regular maize starch (700004337/K-RSTCL; Megazyme International, Ireland) standard extractions performed alongside biological samples.

#### Raffinose Family Oligosaccharide (RFO) quantification

Oligosaccharide extraction was performed as described previously (Dierking and Bilyeu, 2008; Aznar-Moreno et al., 2022). Briefly, sugars were extracted from approximately 15–25 mg of lyophilized tissue using 10 µg/ml ribitol (A5502; Millipore Sigma, Saint Louis, MO, USA) as an internal standard and 1 ml of 50% ethanol. For late developmental and mature seed samples, additional pulverization was performed using a ball mill at 30 Hz for 5 mins or until fully ground. Samples were then heated at 70 °C for 30 mins with intermittent shaking and vortexing. Following heat treatment, samples were centrifuged at 15,000 × g for 15 mins to pellet debris. The supernatant was transferred to a new tube and completely dried using a centrivap. Sugars were reconstituted in 50% methanol and 3 ul sample was separated chromatographically using an Agilent InfinityLab Poroshell 120 HILIC-Z (2.7-µm particle size, 100 × 2.1 mm) column (685775-924; Agilent, Santa Clara, CA, USA). LC-MS/MS data were acquired using a Thermo TSQ Altis mass spectrometer coupled to a Thermo Dionex UltiMate 3000 UHPLC system (Thermo Fisher Scientific, Waltham, MA, USA) and a CTC Analytics HTC-PAL autosampler (CTC Analytics, Zwingen, Switzerland). The LC instrument was operated with a flow of 0.2 ml/min and sugars were eluted using mobile phase buffers 10mM Ammonium formate in water, pH 6.8 (buffer A) and 10mM Ammonium formate (1:9) water: ACN (buffer B) with an oven temperature of 40°C, adapted from prior methods (Czajka et al., 2020). The gradient conditions were as follows (all percentages in v/v): Start (0 min) at 95% B, 3.5 min at 60% B, 6 min at 30% B, 7.5 min at 30% B, 9 min at 95% B, 14.70 min at 95% B. The total run time was 15 min. The TSQ Altis mass spectrometer was operated with a Heated Electrospray ionization (H-ESI) ion source type in negative ionization mode with a static ion spray voltage of -3700 V and a transfer tube temperature of 325°C, sheath/aux/sweep gas settings of 55/7/1 (Arb), and a vaporizer temperature of 350 °C. Direct infusion of standards was performed to determine the diagnostic fragments and collision energies for each sugar to generate the selected reaction monitoring (SRM) approach of targeted sugar determination **(Supplementary materials and methods, Table 1)**. All peaks were integrated using the latest version of Skyline software (22.2-64-bit version; MacCossLab Software, University of Washington, USA)(Pino et al.2020). Quantification was done using a calibration curve.

**Table 1.** Biomass composition of *SDP1-* suppressed soybean seeds. Soybean seed storage biomass was quantified from three to four plants per three reproductive stages and maturity (n=3-4 with 2-4 technical replicates) from WT (Williams 82) and three transgenic genotypes A (*sdp1-1 RNAi A*), B (*sdp1-3, RNAi D*) and C (*sdp1-3, RNAi C*). Total lipid, starch and oligosaccharides (raffinose and stachyose) were determined on a mg per seed basis. Data represents mean ± Standard error. **p≤0.01, and ***p≤0.001, indicate statistically significant difference based on a two-tailed Student’s t-test.

| Developmental Stages | Genotype | Storage reserves (mg/seed) |  |  |  |
|---|---|---|---|---|---|
|  |  | Lipid | Starch | Raffinose | Stachyose |
| R5.5 | WT | 3.91±0.21 | 3.70±0.11 | 0.00±0.00 | 0.00±0.00 |
|  | A | 3.82±0.05 | 2.95±0.08 *** | 0.00±0.00 | 0.00±0.00 |
|  | B | 3.40±0.22 ** | 2.97±0.10 *** | 0.00±0.00 | 0.00±0.00 |
|  | C | 2.95±0.08 *** | 2.39±0.05 *** | 0.00±0.00 | 0.00±0.00 |
| R6 | WT | 19.61±0.37 | 8.72±0.34 | 0.04±0.01 | 0.02±0.01 |
|  | A | 27.68±0.64 *** | 12.26±0.51 *** | 0.05±0.02 | 0.01±0.01 |
|  | B | 33.71±0.36 *** | 13.70±0.83 *** | 0.09±0.03 ** | 0.04±0.03 |
|  | C | 28.24±0.89 *** | 12.43±0.33 *** | 0.09±0.02 ** | 0.02±0.02 |
| R7 | WT | 29.97±0.42 | 3.61±0.27 | 0.87±0.04 | 3.85±0.31 |
|  | A | 41.20±2.96 *** | 3.96±0.76 | 1.11±0.09 *** | 5.90±0.69 *** |
|  | B | 44.36±1.66 *** | 2.32±0.19 *** | 1.50±0.08 *** | 6.85±0.24 *** |
|  | C | 51.80±0.94 *** | 4.03±0.75 | 1.81±0.25 *** | 7.54±0.82 *** |
| R8 | WT | 39.05±0.29 | 0.52±0.05 | 1.48±0.09 | 6.51±0.22 |
|  | A | 52.18±0.65 *** | 0.56±0.02 | 1.70±0.10 ** | 7.41±0.42 *** |
|  | B | 50.78±1.62 *** | 0.47±0.06 | 1.68±0.03 ** | 4.98±0.22 *** |
|  | C | 49.49±0.31 *** | 0.51±0.05 | 1.49±0.04 | 4.22±0.12 *** |

### Transcriptomic analysis of developing soybean seeds

Transcriptomic analysis was performed with seeds from three to four plants per each of *SDP1* RNAi suppressed line B and WT across early (R5.5), mid (R6) and late (R7) developmental stages. Seeds were ground using a GenoGrinder (Cole-Parmer HG-600). Approximately 50 mg of ground tissue was used for total RNA extraction employing the MagMax Plant RNA Isolation Kit in the KingFisher Flex (ThermoFisher Scientific, Waltham, MA, USA) automated system. RNA was quantified using a Nanodrop spectrophotometer and an Agilent TapeStation 4200. Samples with an RNA Integrity Number (RIN) of 8 or higher were deemed of high quality. Library preparation and sequencing were conducted by Novogene using the Illumina NovaSeq X Plus. Briefly, mRNA libraries were prepared with poly(A) enrichment and sequenced using a paired-end 150 bp (PE150) approach. The resulting demultiplexed raw reads were provided in FASTQ format. These were then pseudo-aligned to the WT transcriptome using Salmon v1.10.2 (Patro et al., 2017) default settings. Read count data were imported into R using the tximport package (Soneson et al., 2015). Samples were then defined by their developmental stage (R5.5, R6, R7) and genotype. Differential expression on the integer count values was performed using DESeq2 (Love et al., 2014). Genes with a sum of less than 10 reads or a mean of 20 reads across all samples were excluded from analysis. Principal component analyses (PCA) were performed using the variance stabilized transformed read counts DESeq2. Differential expression analysis was performed using “apeglm”(Zhu et al., 2019) as the Log Fold Change Shrinkage method. Genes were defined as being significantly differentially expressed if they had an adjusted p-value of less than 0.05. Normalized counts were plotted using ggplot and tidyverse (Wickham et al., 2019) functions in R. Gene Ontology (GO) term enrichment analysis was performed using the TopGO R package with the entire set of expressed genes set as background (Alexa et al., 2006). Terms were considered enriched if they passed a p-value of 0.05 on the Fisher test with the “weight01” algorithm and a minimum node size of 100 genes.

### Phosphoenolpyruvate carboxykinase (PEPCK) enzyme activity assay

PEPCK activity assay was performed with developing seeds from four plants, each of WT and three transgenic genotypes. Enzyme extraction and assay procedures were adapted from (Osorio et al., 2014) and (Sharwood et al., 2016). Approximately 100 mg of fresh ground tissue was homogenized in 1 ml of cold extraction buffer containing 50 mM HEPES (pH 7.0), 5 mM dithiothreitol (DTT; freshly prepared), 1% (w/v) polyvinylpolypyrrolidone (PVPP), 2 mM Ethylenediaminetetraacetic acid (EDTA), 2 mM Manganese(II) chloride (MnCl₂), and 0.05% Triton X-100. Samples were vortexed for 30 s, homogenized for 2 min using a bead beater, and incubated on ice for 10 min. The homogenate was centrifuged at 13,000 × g for 30 s at 4 °C, and the clarified supernatant was used for enzyme assays. PEPCK activity was measured in a flat-bottom 96-well microplate. Each reaction contained 10 µL of protein extract and 150 µL of assay buffer comprising 50 mM HEPES (pH 7.0), 4% (w/v) β-mercaptoethanol, 100 mM KCl, 90 mM NaHCO₃, 1 mM ADP, 2 mM MnCl₂, 0.14 mM NADH, 15 mM phosphoenolpyruvate (PEP), and 6 units of malate dehydrogenase. Absorbance at 340 nm was monitored at 25 °C using a Tecan Microplate Reader M 200PRO, with readings taken every 5–10 s for up to 5 min with intermittent shaking. Control reactions lacking PEP were included to correct for background NADH oxidation. PEPCK activity was calculated from the linear decrease in absorbance at 340 nm using the molar extinction coefficient of NADH and expressed as enzyme units (U), defined as µmol NADH oxidized per minute. Protein concentration was determined using the Bradford assay with bovine serum albumin standards, and enzyme activity was normalized accordingly.

### Hormone analysis

#### Phytohormone extraction

Phytohormones including cis-zeatin (cZ), trans-zeatin (tZ), DL-dihyrdrozeatin (DHZ), trans-zeatin riboside (tZR), abscisic acid (ABA), indole-3-acetic acid (IAA), IAA-aspartic acid (IAA-Asp), salicylic acid (SA), 12-oxo-phytodienoic acid (OPDA), jasmonoyl-isoleucine (JA-Ile), and jasmonic acid (JA) were extracted from seeds collected at mid (R6) and late (R7) reproductive stages. Samples were harvested from three to four plants per transgenic genotype and corresponding WT controls. Hormone extraction was performed in triplicate at a tissue concentration of 50 mg ml⁻¹ using ice-cold 1:1 methanol:acetonitrile (MeOH:ACN). For extraction, a stock solvent of 1:1 MeOH:ACN was prepared and supplemented with a mixed stable isotope-labeled internal standard (IS) solution containing 0.4 μM d5-tZ and d5-tZR; 1.0 μM d4-SA, d6-ABA, d5-JA, d5-IAA, ¹³C₆¹ N-IAA-Asp, and d5-dinor-OPDA; and 10.0 μM ¹³C₆¹ N-JA-Ile. The total extraction solvent volume was calculated as (number of samples + 2) × (975 μl MeOH:ACN + 25 μl IS mix). For the experiment described here, 17.55 ml MeOH:ACN and 450 μl IS mix were prepared. Each sample received 1 ml of extraction solvent containing internal standards. Samples were homogenized using a TissueLyser II (Qiagen) for 5 min at 15 Hz, followed by centrifugation at 16,000 × g for 5 min at 4 °C. Supernatants were transferred to fresh 2 ml tubes, and the remaining pellets were re-extracted twice with 800 μl ice-cold 1:1 MeOH:ACN (without IS). All supernatants were pooled and dried using a vacuum centrifuge. Dried extracts were reconstituted in 100 μl of 30% methanol, centrifuged to remove particulates, and filtered through a 0.8 μm polyethersulfone spin filter (Sartorius, Stonehouse, UK). The filtrates were transferred to HPLC vials for subsequent LC–MS/MS analysis. Details of chemicals used for the assay, including reference standards, are included as supplemental material.

#### LC-MS/MS Analysis

Phytohormones including cis-/trans-zeatin (c/tZ), DHZ, tZR, SA, ABA, IAA, IAA-Asp, JA, JA-Ile, and OPDA were quantified using a targeted selected reaction monitoring (SRM) isotope dilution-based LC–MS/MS method **(Supplementary materials and methods, Table 2)**. Quantitative analysis was performed using an Ultimate 3000 UPLC system coupled to a TSQ Altis triple quadrupole mass spectrometer equipped with a heated electrospray ionization (HESI) source (Thermo Fisher Scientific, Waltham, MA, USA). A 5 μl aliquot of each reconstituted sample was injected onto a 3.0 × 100 mm, 1.8 μm ZORBAX Eclipse XDB-C18 column (Agilent Technologies, Santa Clara, CA, USA). Chromatographic separation was achieved over 22 min using a binary gradient consisting of mobile phase A (0.1% acetic acid in water) and mobile phase B (0.1% acetic acid in 3:1 acetonitrile:methanol). The gradient program was as follows: 10% B from 0 to 2.0 min, increased to 25% B at 7.0 min, ramped to 65% B at 15.0 min, increased to 95% B at 18.0 min, and held at 95% B until 24.0 min. The flow rate was maintained at 0.3 ml min⁻¹.Mass spectrometry source parameters were set as follows: sheath gas, 60 arbitrary units; auxiliary gas, 5; sweep gas, 2; ion transfer tube temperature, 325 °C; vaporizer temperature, 350 °C; and spray voltage of +3000 V in positive ion mode and −3300 V in negative ion mode. MS conditions included a collision-induced dissociation (CID) gas pressure of 1.5 mTorr, with Q1 and Q3 resolutions set to 0.7 full width at half maximum (FWHM) and a cycle time of 0.6 s. Analytes and corresponding internal standards were monitored using optimized SRM transitions in a polarity-switching acquisition method, with cytokinins and auxins detected in positive ion mode and other analytes detected in negative ion mode. Data acquisition was performed using Xcalibur software (version 4.2.47; Thermo Fisher Scientific), and quantitative analysis was conducted using QuanBrowser (Thermo Fisher Scientific). Phytohormone concentrations were determined by comparing analyte-to-internal standard peak area ratios to calibration curves generated using matched standards and internal standards. Internal standard concentrations in calibration solutions were 2.5 μM for ¹³C₆¹ N-JA-Ile, 0.10 μM for d5-tZ and d5-tZR, and 0.25 μM for all other internal standards. Mixed calibration standards were prepared over a range of 0.5 fmol to 250 pmol on-column, and the final calibration range for each analyte was adjusted based on its abundance in the samples.

**Table 2.**
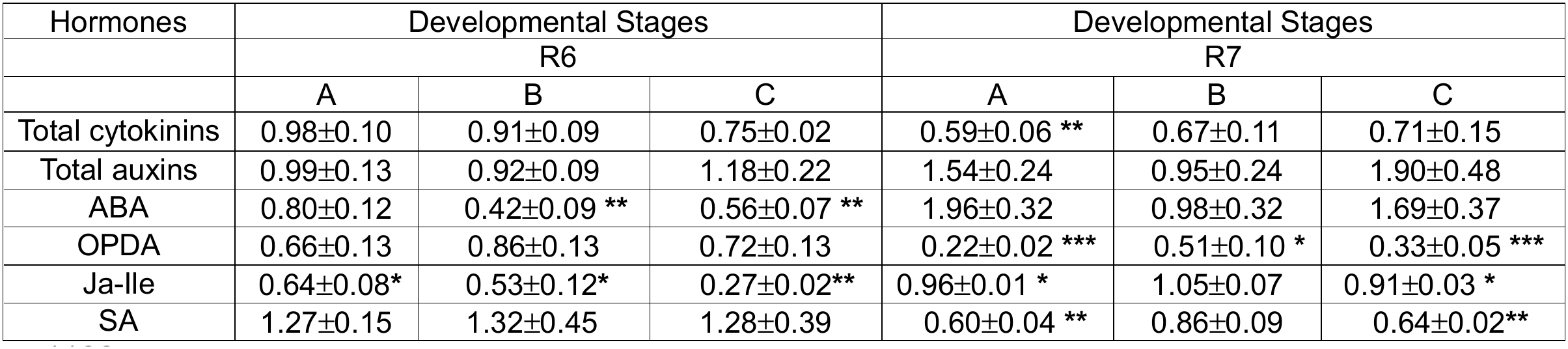
Relative levels of total cytokinins, total auxins, abscisic acid (ABA), 12-oxo-phytodecanoic acid (OPDA), Jasmonoyl-isoleucine (Ja-Ile), and salicylic acid (SA), in the three *sdp1* suppressed transgenic genotypes A (*sdp1-1 RNAi A*), B (*sdp1-3, RNAi D*) and C (*sdp1-3, RNAi C*) as compared to WT during R6 (mid) and R7 (late) development stages. Data represents mean ± standard error (n=3-4 plants/genotype). *p ≤ 0.05, **p ≤ 0.01, ***p ≤ 0.001 indicates a statistically significant difference based on a two-tailed t-test.

| Hormones | Developmental Stages |  |  | Developmental Stages |  |  |
|---|---|---|---|---|---|---|
|  | R6 |  |  | R7 |  |  |
|  | A | B | C | A | B | C |
| Total cytokinins | 0.98±0.10 | 0.91±0.09 | 0.75±0.02 | 0.59±0.06 ** | 0.67±0.11 | 0.71±0.15 |
| Total auxins | 0.99±0.13 | 0.92±0.09 | 1.18±0.22 | 1.54±0.24 | 0.95±0.24 | 1.90±0.48 |
| ABA | 0.80±0.12 | 0.42±0.09 ** | 0.56±0.07 ** | 1.96±0.32 | 0.98±0.32 | 1.69±0.37 |
| OPDA | 0.66±0.13 | 0.86±0.13 | 0.72±0.13 | 0.22±0.02 *** | 0.51±0.10 * | 0.33±0.05 *** |
| Ja-Ile | 0.64±0.08* | 0.53±0.12* | 0.27±0.02** | 0.96±0.01 * | 1.05±0.07 | 0.91±0.03 * |
| SA | 1.27±0.15 | 1.32±0.45 | 1.28±0.39 | 0.60±0.04 ** | 0.86±0.09 | 0.64±0.02** |

[illegible]

### Lignocellulosic wall isolation

Lignocellulosic wall content was extracted from plants each of WT and the three RNAi lipase suppressed genotypes A, B and C. Alcohol Insoluble Residue (AIR), as described earlier (Caffall et al., 2009; Foster et al., 2010; Pettolino et al., 2012; Kataya et al., 2025) was used for the extraction of lignocellulosic wall material. Briefly, ∼100 mg of powdered lyophilized seed samples was sequentially extracted with 80% ethanol (v/v), 1:1 (v/v) chloroform: methanol, 100% methanol and 100% acetone. The pellet was resuspended in each solvent and incubated for 10 mins at room temperature on a rotating mixer. Between each extraction, samples were pelleted by centrifugation at 10,000 g for 10 mins using a table-top microcentrifuge. After solvent extraction, the pellet was washed twice with water and subsequently used for starch removal. Mature soybean seeds contain minimal starch (Kambhampati et al., 2021). However, to prevent interference from trace amount of starch in the sample, alpha-amylase starch digestion was performed. The resulting pellet was washed with sterile water and dried overnight at 40°C using a centrifugal vaccum concentrator. The dried AIR material represented the lignocellulosic (cell wall) enriched fraction.

### Statistical Analysis

To test for a significant difference of the means of different parameters between the WT and the *SDP1*-suppressed genotypes, two-tailed Student’s t-test of unequal variance was performed. Means were considered significantly different at p≤0.05. Pearson correlation analysis was performed, and p-value was determined using the corresponding t-distribution of the Pearson correlation coefficient (PCC) to determine significance of the correlation analysis.

## Results

### Seed-specific suppression of SDP1 enhances seed weight and seed area

Previously, we found that suppressing the *SDP1* gene increases seed weight (Aznar-Moreno et al., 2022), consistent with earlier observations in Arabidopsis (Kim et al., 2014) and soybean (Kanai et al., 2019). In this study, we conducted a detailed greenhouse experiment to characterize the increase in seed size during development. Four to six plants were grown from three independent homozygous *SDP1* suppressed lines A, B, and C, with the WT controls. As vegetative growth influences the canopy and thus the productivity from a plant; node number, pod number, total seeds were measured from each of the individual T4 plants per line and WT controls. No significant difference was obtained in node, pod, or seed number relative to the control plants **(Figure 1A-C).** Next, the 100-seed weight was assessed in each plant. The results indicated that seeds from the lipase suppressed transgenic soybean genotypes had significantly higher mass than the WT (p ≤ 0.001), resulting in an 18-25% increased seed weight **(Figure 1D and Supplementary figure S1C)**. Pod and node number were modestly correlated (r=0.52, p=0.06; **Supplementary figure S1A)**. Seed and pod number were strongly related (r=0.88, p ≤ 0.001; **Supplementary figure S1B**); however, seed weight did not correlate with seed number (r=-0.25, p=0.647) **(Figure 1E)**. Seed weight was also compared to seed area. Seed area increased by 13-21% (p ≤ 0.001), compared to the control **(Figure 1F and Supplementary figure S1D)**. The results indicated that *SDP1* suppression in soybean seeds did not affect plant architecture and branching apart from the change in seed size. As the change in seed weight was not associated with seed number, total seed biomass was increased on a per plant basis.

**Figure 1.**
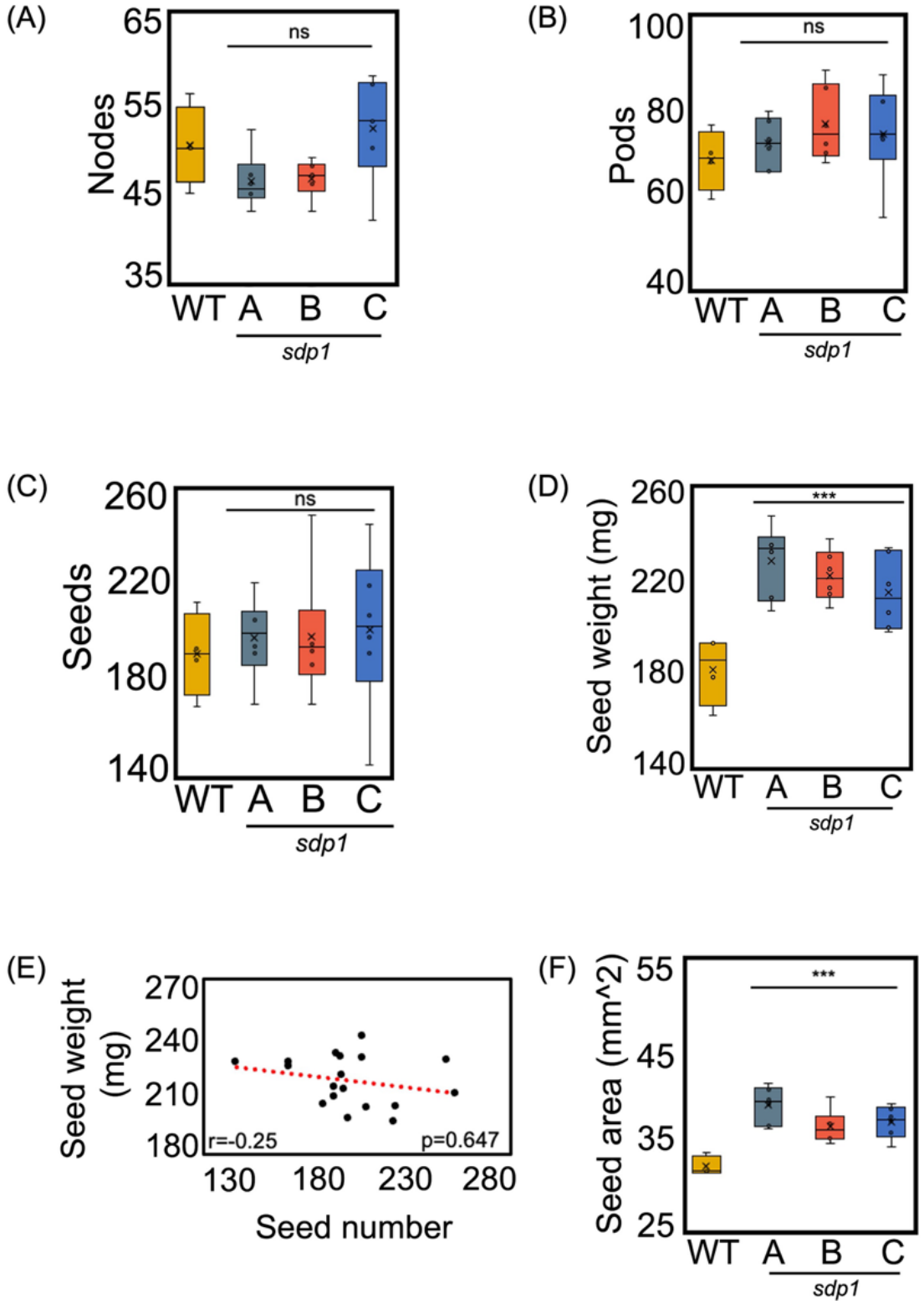
Phenotypic and agronomic measurements of WT and *SDP1-* suppressed soybean genotypes The total number of nodes (A), pods (B), and seeds (C) were determined from four-six plants each for WT, A (*sdp1-1 RNAi A*), B (*sdp1-3, RNAi D*) and C (*sdp1-3, RNAi C*), labeled here as *sdp1.* Average individual seed weight was calculated from 100 seeds per each of four-six plants per genotype (D). The correlation between average individual seed weight and seed number (E) was determined using the Pearson correlation coefficient (PCC) and the p-value was determined using the corresponding t-distribution of the PCC. Average individual seed area was measured using RGB images of 10 seeds from each of four-six plants per genotype, processed with PlantCV software (F). Box-plot center lines indicate the median, ‘x’ represent the mean; box limits, 25^th^ and 75^th^ percentiles; whiskers, 1.5 times the interquartile range from the 25^th^ and 75^th^ percentiles. ***p ≤ 0.001 indicates a statistically significant difference based on a two-tailed Student’s t-test.

### Enhanced seed size is due to increased cell size with no significant change in cell number

We sought to understand if the significant increase in seed size was attributable to changes in cell number or morphology. We examined seed cross sections for each of the three RNAi genotypes and compared them to the WT. Cells of all *SDP1-*suppressed mature seeds showed a significant increase in cell area relative to the WT, with cells of lines A, B and C showing 11, 12 and 4% increased area, respectively (p ≤ 0.001; **Figure 2A-E)**. In contrast, no significant change in cell number was observed **(Figure 2F)**.

**Figure 2.**
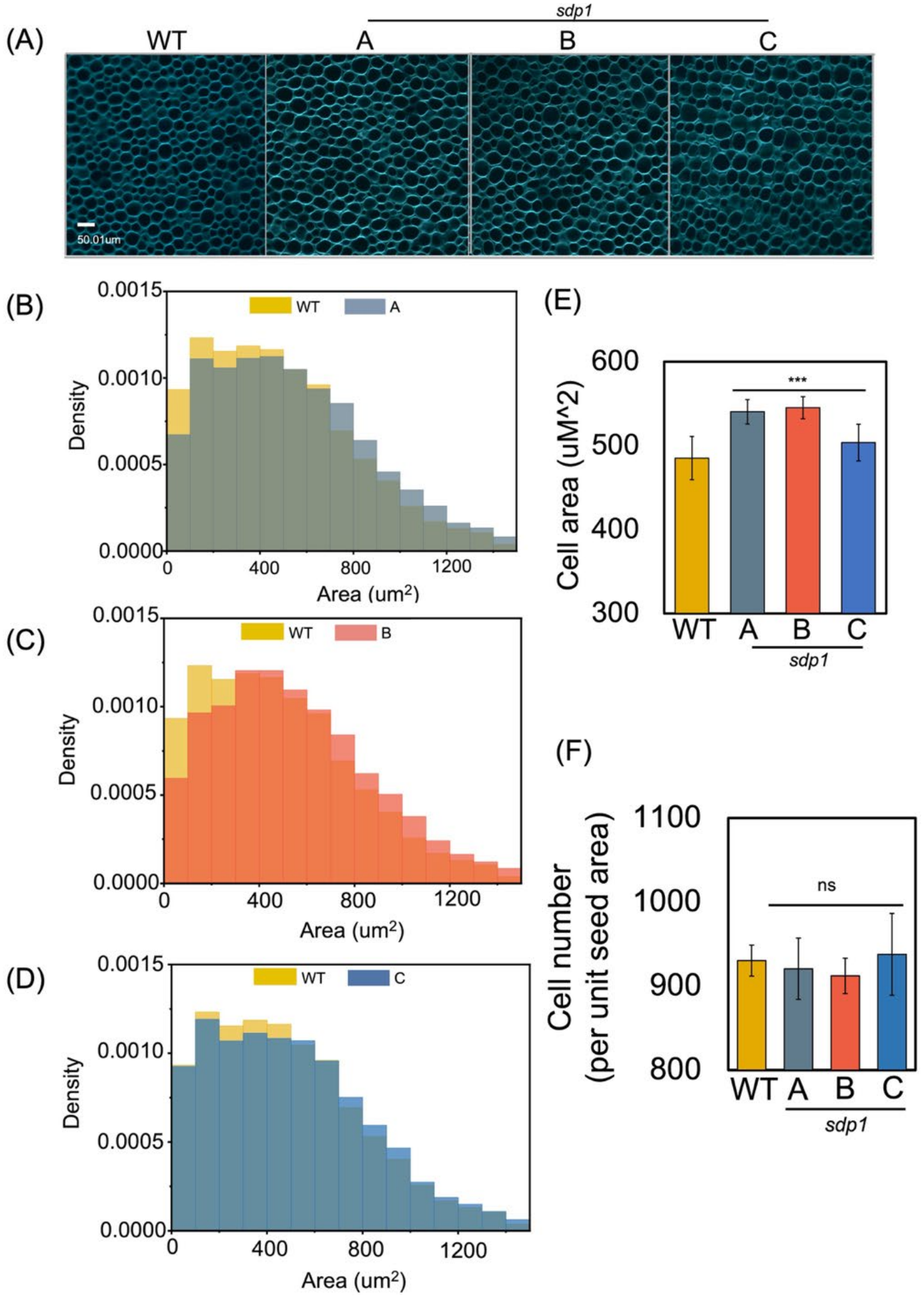
Cellular morphology of WT and *SDP1-* suppressed soybean genotypes Confocal microscopy imaging of three to five matured soybean seeds from each of WT and three *sdp1* suppressed genotypes A (*sdp1-1 RNAi A*), B (*sdp1-3, RNAi D*) and C (*sdp1-3, RNAi C*), labeled here as *sdp1*, bar=50.01 um (A) Density histogram using 4-10 images per seed from three to five seeds per each of WT and A (B) WT and B (C) and WT and C (D) indicated a shift towards bigger cell area in the transgenic seeds as compared to WT controls. Cell area (E) and cell number (F) of *SDP1*-suppressed and WT genotypes (F). Data represents mean ± standard error. ***p ≤ 0.001 indicates a statistically significant difference based on a two-tailed Student’s t-test.

### RNAi sdp1 seed weight is increased by mid-development

The *sdp1* mutation enhanced soybean seed size at maturity. WT and SDP1 altered seeds matured at the same time; however, it was unclear when larger seeds emerged during seed filling. Fresh weight (FW) and dry weight (DW) were analyzed from developing WT and lipase-altered genotypes, harvesting seeds together. During the early stages of filling (R5.5), lines A, B and C did not differ relative to WT **(Figure 3)**; however, by mid development (R6) and through later stages (R7), differences could be observed. A, B, and C were 33%, 48% and 38% greater in FW and 43%, 61% and 45% increase in DW compared to WT at R6. At R7, the same lines had 33%, 33% and 51% greater FW and 35%, 32% and 57% higher DW respectively.

**Figure 3.**
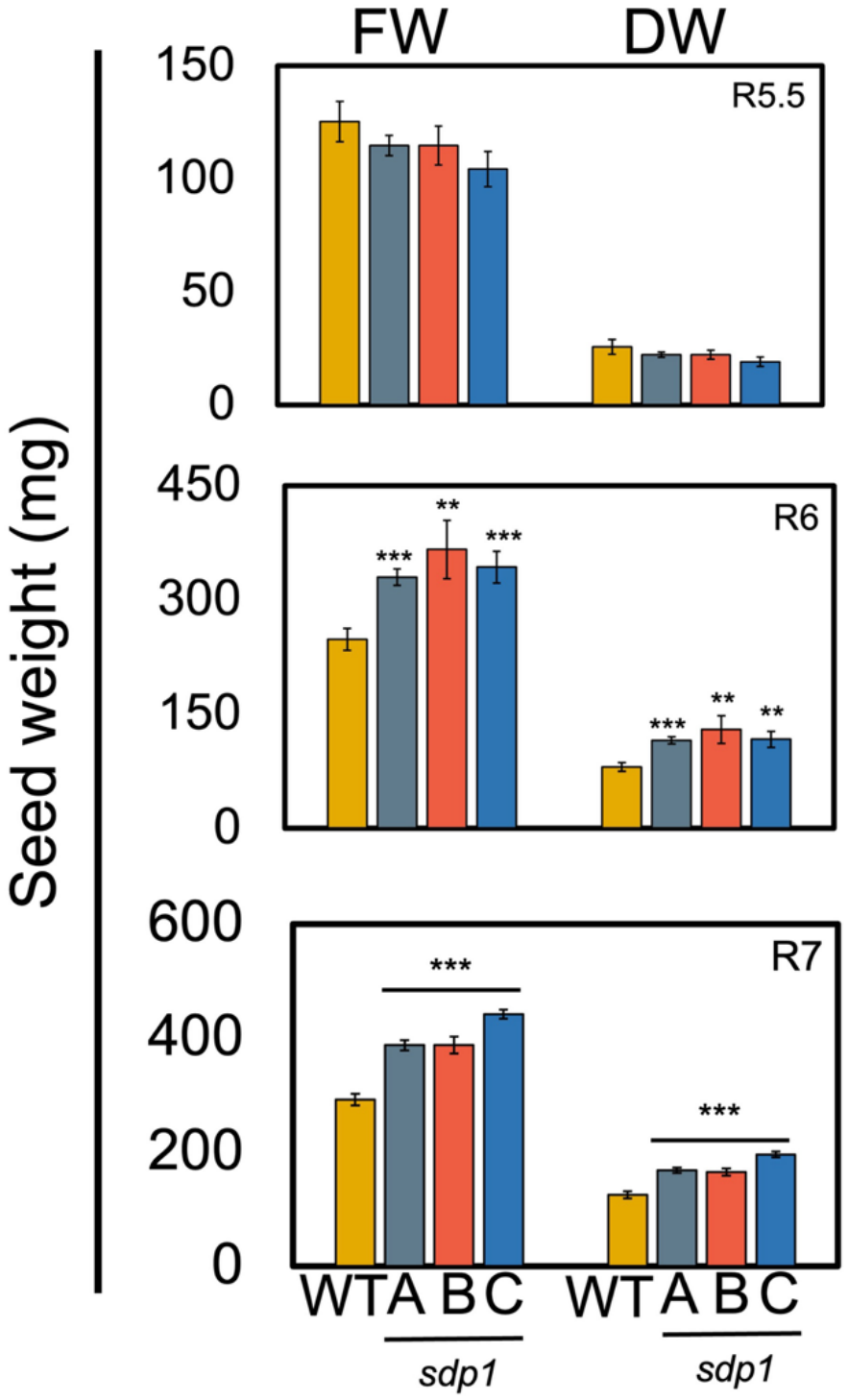
Fresh weight and dry weight measurement of soybean seeds Fresh weight (FW) and dry weights (DW) measurements of seeds from four individual plants of the three *SDP1-* suppressed genotypes A (*sdp1-1 RNAi A*), B (*sdp1-3, RNAi D*) and C (*sdp1-3, RNAi C*), labeled here as *sdp1* and WT. Data represents mean ± standard error. **p ≤ 0.01 ***p ≤ 0.001 indicates a statistically significant difference based on a two-tailed Student’s t-test.

### Developmental composition of soybean seed biomass indicated elevated oil levels

The contributions to altered single seed weight **(Figure 1 D)** were further evaluated by assessing seed composition over development. T4 seeds were collected at R5.5, R6, R7 development stages, and at maturity (R8). Biomass was analyzed from three to four plants, with two to four technical replicates per plant, for each transgenic line and WT. Biomass analysis on a per seed basis was obtained from percent biomass storage reserve content **(Supplementary table S1)** and average individual seed weight **(Figure 1 D and Figure 3).**

FAME analysis indicated that at R5.5, total lipid content (mg/seed) was reduced in all *SDP1*-suppressed lines compared to WT. This difference was significant for B (p ≤ 0.01) and C (p ≤ 0.001). From R6 through R8, oil content was elevated in all three transgenic lines compared to WT (p ≤ 0.001) **(Table 1).** The fatty acid profile of the transgenics changed over development, specifically in the levels of palmitic (C16:0) and linoleic (C18:2) acid. Stearic acid (C18:0) was increased at R5.5 for B (p ≤ 0.01) and C (p ≤ 0.001) and the difference decreased until maturity. Oleic acid (C18:1) was elevated for A (p ≤ 0.05, p ≤ 0.01 and p ≤ 0.001). Linolenic acid (C18:3) on the other hand significantly reduced for selected transgenic genotype (p ≤ 0.05 and p ≤ 0.001) across development respectively **(Supplementary table S2)**.

Starch content in soybeans is transient and peaks at R6 and declines through R7 and R8 (Kambhampati et al., 2021). This trend was observed in both WT and *SDP1*-suppressed seeds **(Table 1);** however, at R5.5, transgenic seeds had significantly lower starch than WT (p ≤ 0.001). The starch levels in transgenics increased at R6, matched WT levels by R7 in most lines **(Table 1)** and were similar at maturity.

Mature soybeans contain appreciable quantities of RFOs (Kambhampati et al., 2021) including raffinose and stachyose that cannot be digested by monogastric animals and detract from the overall value of soybean. Prior studies indicated that *GmSDP1* suppression reduces the level of RFOs, particularly stachyose, on a percent biomass basis at maturity (Aznar-Moreno et al., 2022). Here, the RFOs in *SDP1*-suppressed soybean genotypes were compared to the WT over development. Raffinose and stachyose were undetected in R5.5 seeds **(Table 1)**. Mid-development (R6) seeds had elevated raffinose in B and C (p ≤ 0.01) **(Table 1)** without changes in stachyose. R7 seeds were elevated in raffinose and stachyose levels in A, B, and C seeds (p ≤ 0.001) **(Table 1)**. Consistent with prior observations at maturity (Aznar-Moreno et al., 2022), our results indicated a decline in raffinose and stachyose levels on a percent biomass basis in the T4 mature seeds of *SDP1*-suppressed lines **(Supplementary table S1)**; however, due to enhanced seed size, total raffinose content (mg/seed) of mature *SDP1*-suppressed seeds was higher than WT controls, with the difference being statistically significant for A and B (p ≤ 0.01) **(Table 1)**. Stachyose levels in lines B and C (p ≤ 0.001) were reduced relative to the WT, except for A, which showed an increase in stachyose content (p ≤ 0.001) **(Table 1)**. The metabolic consequences of these changes were evident in the mature seeds. The absolute amount of oil produced was increased by 27% to 34% per seed, because of the larger seed size and the proportional increase in oil (**Supplementary table S3)**. Total raffinose was elevated 1% to 15% (**Supplementary table S3)**; however, starch and stachyose changes were genotype-specific (**Supplementary table S3).**

### Increased number of differentially expressed genes during late development

To better understand the mechanism by which silencing *SDP1* enhances cell size **(Figure 2A-E)** and increases seed size **(Figure 1D and 1F),** we performed RNAseq across seed development. Principal Component Analysis (PCA) of the transcriptomic data from the WT and *SDP1*-suppressed RNAi samples from line B showed consistent clustering between developmental timepoints, with minimal differences between WT and RNAi samples. PCA indicated that the first principal component (PC1) accounted for 88% of variance separating samples from early (R5.5 and R6) and late (R7) seed development. The second principal component (PC2) with 5% variance reflected smaller changes associated with changes between early (R5.5) and mid-stage (R6) development **(Figure 4A)**.

**Figure 4.**
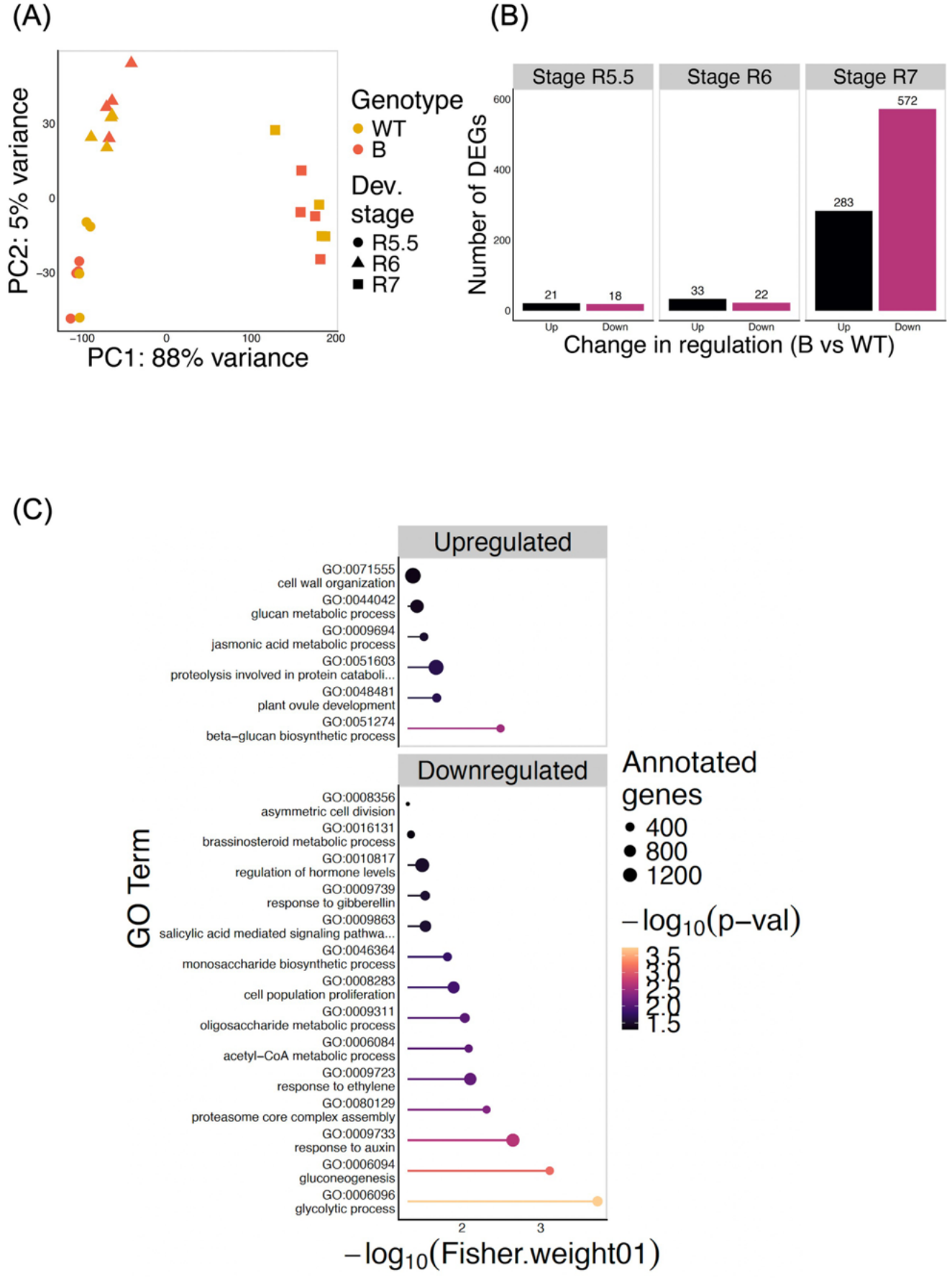
Stage specific differential transcriptomic responses to *SDP1* suppression Soybean seeds from three to four plants per three reproductive stages (R5.5, R6 and R7) from WT and the transgenic genotype B (*sdp1-3, RNAi D*), was used to determine transcriptomic responses to *SDP1* suppression. Principal component analysis (PCA) (A) Number of differentially expressed genes (DEGs) (B) and Selected Gene Ontology (GO) term enrichment analysis of R7 seeds (C). The colors of the lines represent the –log10(Fisher.weight01) p-value as calculated by the topGO R package.

When comparing gene expression between RNAi lines and the WT control, R7 stage contained the highest number of differentially expressed genes (DEGs) with 283 genes and 572 genes being up and down regulated respectively. In contrast, very few DEGs were identified in R5.5 and R6 **(Figure 4B and Supplemental file S1)**. As the R7 stage showed the highest number of DEGs, we performed a Gene Ontology (GO) enrichment analysis at this stage. Downregulated genes were significantly enriched for various metabolic processes including hormone associated metabolic processes, acetyl-CoA metabolic processes, oligosaccharide metabolic processes, monosaccharide biosynthetic process, gluconeogenesis, glycolytic processes, proteasome core complex assembly processes and metabolic processes associated with cell division and proliferation **(Figure 4C).** Selected GO term enrichment of upregulated genes was associated with cell wall organization including glucan metabolic processes, beta-glucan biosynthetic processes, genes associated with JA metabolic processes and plant ovule development **(Figure 4C).** A detailed list of all DEGs and GO-term enrichment are included in **Supplemental file S1.**

### Seed-specific SDP1 suppression indicated altered expression of candidate genes from signaling pathways to regulate seed size

A closer inspection of the developmental dataset identified candidate genes of the F-Box family proteins and Really Interesting New Gene (RING)/U-Box super family proteins as differentially expressed between genotypes. F-box and RING/U-box superfamily proteins contain members that are essential components of the ubiquitin-proteasome system and are known to regulate seed size through interactions with phytohormones like JA signaling pathways (Koo et al., 2014; Durand et al., 2016; Hu et al., 2021). At the late developmental stage (R7), expression of five F-box family genes and RING/U-box genes was significantly altered **(Supplementary figure S2, S3).** Further, two members of the Protein Phosphatase 2C (PP2C) family proteins were also differentially regulated over development **(Supplementary figure S4)**.

### Reduced PEPCK activity supports suppressed gluconeogenesis in RNAi genotypes

Isotopic labeling studies suggest that gluconeogenesis may occur late in seed maturation when no phloem-based source of sugars is available (Kambhampati et al., 2021). The *SDP1* suppressed transgenic seeds indicated downregulated genes associated with gluconeogenesis **(Figure 4C and Supplemental file S1).** Although changes in transcript levels are not indicative of altered flux, a reduction in gluconeogenesis would be consistent with reduced lipid turnover and fatty acid breakdown.

To further evaluate this, the activity of PEPCK, which is considered a rate-limiting enzyme for gluconeogenesis, was analyzed over the course of soybean seed development in the three transgenic genotypes and WT **(Figure 5)**. R5.5 seeds of *sdp1* RNAi line A, B and C had 16-41% of PEPCK activity measured in WT. Similarly, R6 and R7 seeds were reduced relative to WT. PEPCK activity was highest in the R7 stage. The differences in PEPCK activity between the transgenic soybean seeds and the WT seeds were statistically significant at all developmental stages except for B at R6 stage **(Figure 5)** and were consistent with a reduction in gluconeogenic capacity that may be important when phloem sugars are not available. Transcripts for several related genes that would be consistent with a role for gluconeogenesis, including malate dehydrogenase, pyruvate kinase, and pyruvate dehydrogenase, were reduced at R7 in the SDP1-suppressed seeds (**Supplemental file S1)** consistent with a reduction in this pathway late in development.

**Figure 5.**
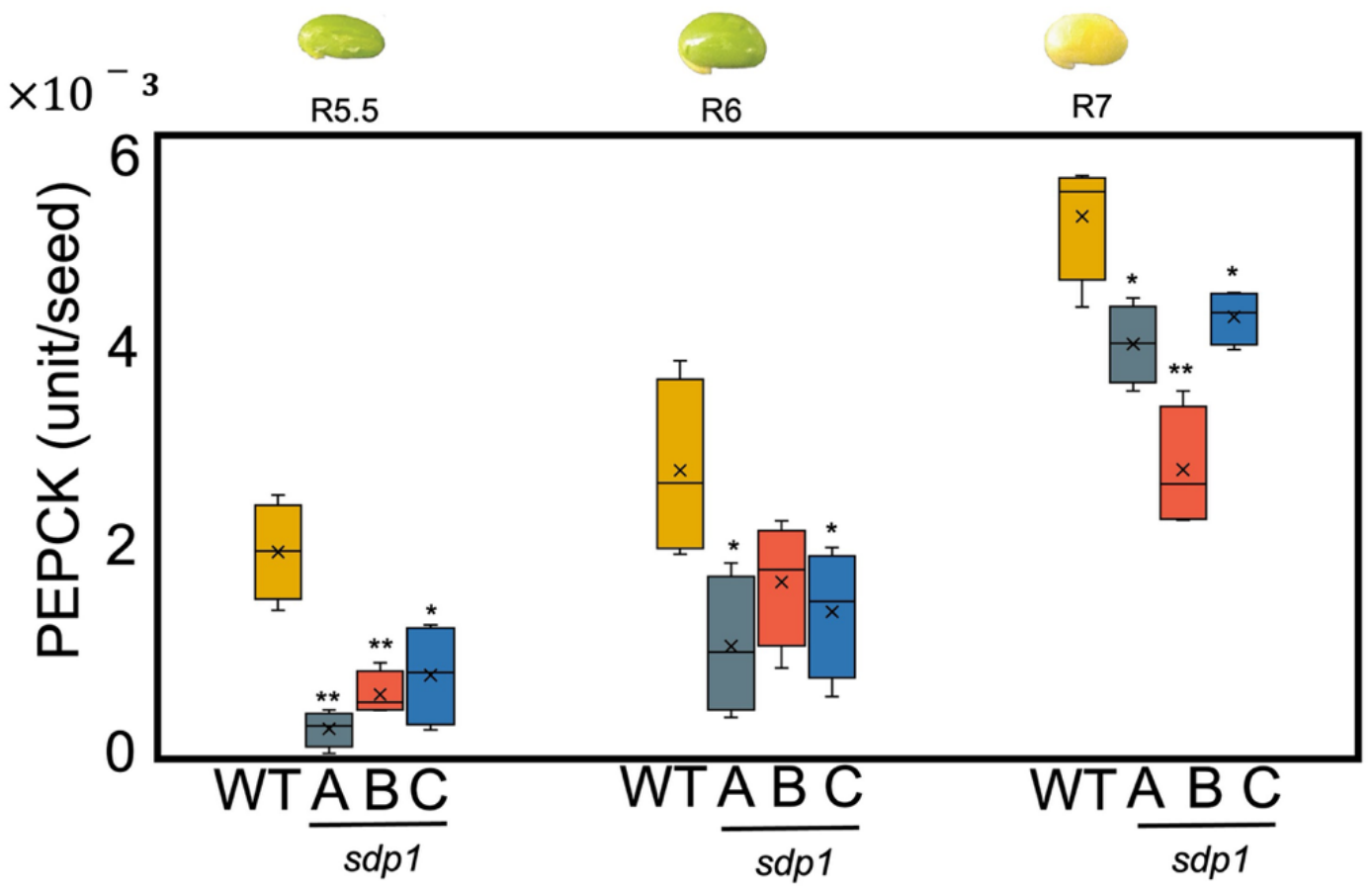
PEPCK activity across soybean seed development PEPCK activity, as a signature of gluconeogenesis, over the course of seed development from four individual plants of the three *SDP1-* suppressed genotypes A (*sdp1-1 RNAi A*), B (*sdp1-3, RNAi D*) and C (*sdp1-3, RNAi C*), labeled here as *sdp1* and WT. Box-plot center lines indicate the median, ‘x’ represent the mean; box limits, 25^th^ and 75^th^ percentiles; whiskers, 1.5 times the interquartile range from the 25^th^ and 75^th^ percentiles. *p ≤ 0.05, **p ≤ 0.01 indicates a statistically significant difference based on a two-tailed Student’s t-test.

### SDP1 suppression results in altered phytohormone levels during mid- and late-stage seed development

Gene expression values suggested that seed-specific suppression of *SDP1* might play a role in altering the levels of genes associated with hormone regulation and hormone metabolic processes during late seed development **(Figure 4C and Supplemental file S1)**. Phytohormones, including cytokinins, auxin, ABA, and JA, are major regulators of seed size and morphology (Cao et al., 2020; Hu et al., 2021; Sharma et al., 2022; Zhang et al., 2025a; Zhang et al., 2025b). Some of them control the turnover of key transcription factors through interactions with F-box protein-mediated ubiquitination, thereby integrating hormonal regulation with the balance between plant growth and defense (Santner and Estelle, 2010; Durand et al., 2016). To evaluate the impact of *SDP1* suppression on phytohormone homeostasis, targeted LC–MS/MS analysis was performed to quantify total cytokinins (sum of tZ, cZ, DHZ, and tZR), total auxins ( IAA, IAA-Asp), ABA, SA, OPDA, JA, and JA-Ile at two stages of seed development: R6 and R7. JA was not detected at either developmental stage. Relative levels of hormones for the three *sdp1* RNAi lines were compared to WT (**Table 2)** with absolute levels included in **Supplemental file S2.**

Notably, OPDA was reduced in all *sdp1* RNAi lines at both developmental stages, with more pronounced decreases at R7. At R6, OPDA levels declined by 34%, 14%, and 28% in A, B, and C, respectively. At R7, reductions were 22-51% of WT **(Table 2)**. OPDA is the precursor of JA biosynthesis and plays an important role in balancing the growth-defense tradeoff in crops (Zhu et al., 2026). Other components of the JA signaling pathway provided further supporting evidence of the role of JA in regulating seed yield. JA-Ile levels were decreased in the transgenic lines, at R6 and in two out of the three transgenic genotypes at R7 **(Table 2)**.

Suppression of JA-Ile can positively regulate seed size (Hu et al., 2021; Jiang et al., 2025; Zhang et al., 2025a). Thus, changes in OPDA and JA-Ile support an SDP1 mechanism that affects F-box and RING-based ubiquitination of growth-related transcription factors, including those associated with seed size.

Other hormones were altered by the *SDP1* suppression. ABA is a major regulator of seed development, controlling seed maturation, storage reserve accumulation, dormancy, and germination. In Arabidopsis, ABA levels peak at distinct times during early and late seed development (Zhang et al., 2025b). Consistent with this dynamic regulation, ABA content in *SDP1*-suppressed lines differed from WT in a stage-dependent manner. During R6, ABA content was significantly reduced in lines B and C by 58% or 44%, respectively, relative to WT. The reduction in line A was moderate (20% relative to WT). By R7, ABA levels were increased in lines A and C **(Table 2)**.

Total cytokinin levels were lower in all transgenic lines relative to WT at all stages (Table 2). Cytokinins are well known for their roles in regulating cell division and seed development, reviewed in(Sharma et al., 2022), but their effects on plant growth and development are complex and context-dependent, with contrasting effects reported during root and shoot meristem development (Lindsay et al., 2006; Werner et al., 2010; Bianco et al., 2013). The larger-seed phenotype observed in the *SDP1*-suppressed genotype, despite the absence of detectable changes in seed number **(Figure 1E)** or cell number **(Figures 2F and 4C)** and the reduction in cytokinin levels, suggests that the relationship between cytokinin homeostasis and seed size is not straightforward

Auxin and SA effects were variable and not easily interpreted **(Table 2)**. Auxin regulates seed development through its effects on silique, embryo, endosperm, and seed coat development (reviewed in (Cao et al., 2020). Previous studies have shown that auxin accumulation and distribution vary throughout seed development (Lur’ and Setter, 1993; Bernardi et al., 2012; Locascio et al., 2014). Similarly, we found that total auxin levels exhibited genotype- and stage-specific variation that was more difficult to clearly assess **(Table 2).** SA levels showed an apparent stage-dependent response to *SDP1* suppression (**Table 2)** but the relationship with seed size remains unclear.

### Seed-specific suppression of SDP1 enhances lignocellulosic cell wall content

Suppression of *SDP1* resulted in a significant increase in soybean seed size **(Figure 1D and 1F)**. Microscopy analysis revealed that this phenotype was primarily driven by an increase in cell area **(Figure 2E)**, with no detectable change in cell number **(Figure 2F)**. Based on these observations, we hypothesized that enlarged cell size would be accompanied by alterations in cell wall composition in the RNAi lipase lines. Consistent with this hypothesis, transcriptomic analysis of late-developing (R7 stage) soybean seeds showed upregulated genes involved in the β-glucan biosynthetic process **(Figure 4C and Supplemental file S1)**. To further investigate this, we quantified total lignocellulosic content in mature (R8 stage) seeds from the three independent transgenic lines—A, B, and C—alongside WT controls. All transgenic lines exhibited a significant increase in total lignocellulosic content per seed compared to the WT based on per seed **(Figure 6)**. Specifically, transgenic line C increased by 23% (p ≤ 0.05) relative to WT, A was elevated. by 30% (p ≤ 0.01), and B by 32% (p ≤ 0.001) **(Figure 6; Supplemental file S3).**

**Figure 6.**
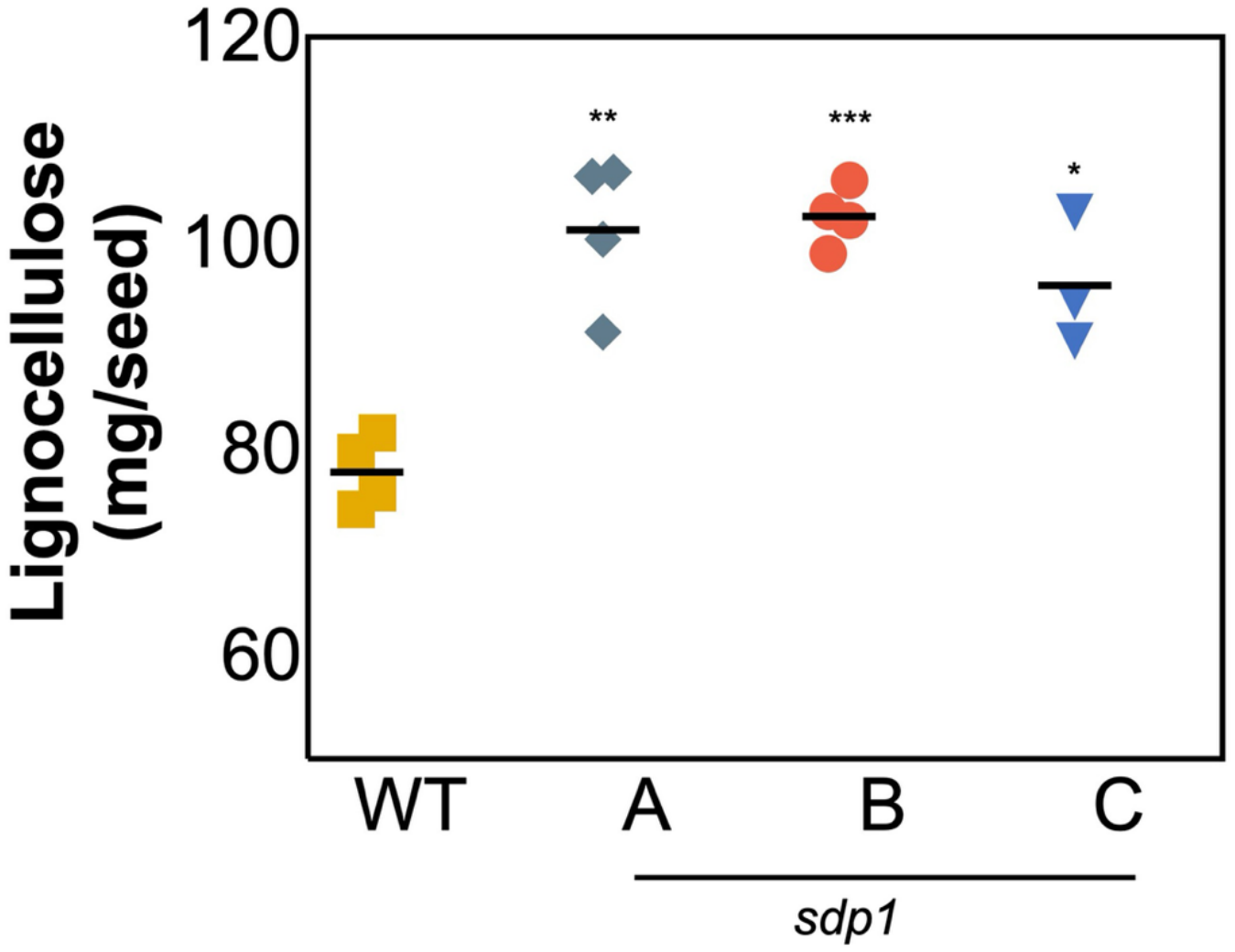
Lignocellulose content Lignocellulose wall content from four plants each of the three *SDP1-* suppressed genotypes A (*sdp1-1 RNAi A*), B (*sdp1-3, RNAi D*) and C (*sdp1-3, RNAi C*), labeled here as *sdp1* and WT. Data represents mean ± standard error. *p ≤ 0.05, **p ≤ 0.01, ***p ≤ 0.001 indicates a statistically significant difference based on a two-tailed Student’s t-test.

## Discussion

### SDP1 suppression enhances soybean seed size through increased cell size

Seed size is a key agronomic trait that directly influences crop fitness and yield (Moles et al., 2005; Fan et al., 2006; Song et al., 2007; Linkies et al., 2010; Li and Li, 2014; Zhang et al., 2024a; Zhang et al., 2024 b; Wang et al., 2025; Zhang et al., 2025b) Mature seed size is determined by the coordinated development of three genetically distinct compartments—the maternally derived seed coat, the diploid embryo, and the triploid endosperm (Lopes and Larkins’, 1993; Reiser and Fischer, 1993; Chaudhury and Berger, 2001; Haughn and Chaudhury, 2005; Zhang et al., 2025a). Interconnected networks of gene expression, phytohormone signaling, nutrient availability, and other signaling pathways collectively perform developmental regulation and control embryo and endosperm growth and ultimately influence seed yield traits (Garcia et al., 2005; Zhang et al., 2025a). Relatively few studies have identified genes that simultaneously improve both seed oil accumulation and yield. Our results support a conserved role for *SDP1* suppression in promoting increased seed weight that is sustained through seed filling to maturity **(Figure 1D, F and Figure 3)**. These findings further strengthen the potential of seed-specific *SDP1* suppression as a biotechnological strategy to improve both seed yield and oil content (Kanai et al., 2019; Aznar-Moreno et al., 2022), without compromising protein accumulation (Aznar-Moreno et al., 2022). By combining of microscopy, transcriptomic, and biochemical analyses, we provide mechanistic insight into the phenotype. Changes in seed size were the result of increased cell area **(Figure 2E)**, indicating that cell expansion, rather than enhanced cell proliferation, underlies the observed phenotype. This interpretation is supported by our transcriptomic analyses, which revealed downregulated genes associated with cell division and upregulated genes involved in cell wall organization and β-glucan biosynthetic processes **(Figure 4C; Supplemental file S1)**. Furthermore, the significant increase in lignocellulosic cell wall content **(Figure 6)** provided additional physiological evidence supporting enhanced cell wall remodeling during seed development.

### Enhanced seed size is associated with altered JA signaling

The seed-specific suppression of *SDP1* was associated with altered JA-related hormone profiles during soybean seed development. Both OPDA and JA-Ile were significantly reduced in *SDP1*-suppressed seeds relative to WT **(Table 2)**. OPDA is a precursor in the JA biosynthetic pathway (Tien et al., 2013), and JA signaling has established roles in regulating plant growth and development (Tien et al., 2013; Hu et al., 2021; Jiang et al., 2025; Kamble, 2025). Although the effects of JA on growth can vary among species and developmental context (Hyun et al., 2008; Singh et al., 2011), the concurrent reduction in OPDA and JA-Ile is consistent with altered JA metabolism in *SDP1*-suppressed seeds. Because JA-Ile is the bioactive conjugate that promotes JA signaling, reduced accumulation raises the possibility that JA signaling is attenuated in *SDP1*-suppressed seeds.

Transcriptomic analysis showed upregulated genes associated with JA metabolic processes and plant ovule development **(Figure 4C; Supplemental file S1)**. These transcriptional changes, together with the reduced accumulation of OPDA and JA-Ile, provide independent evidence of altered JA-associated processes during seed development. Previous genetic studies provide support for a relationship between reduced JA signaling and increased seed or reproductive organ size (Hu et al., 2021; Wang et al., 2023; Zhang et al., 2025a). Mutant analyses of positive and negative regulators of JA signaling have demonstrated effects on seed development and size (Hu et al., 2021; Wang et al., 2023; Zhang et al., 2025a). Thus, a reduction in JA-Ile, along with altered expression of genes associated with plant ovule development and JA metabolism support the possibility that altered JA signaling contributes to the enhanced seed size phenotype observed following *SDP1* suppression.

The transcriptomic analysis also identified changes in components of the ubiquitin-mediated protein degradation system. GO annotation analysis revealed downregulation of genes associated with proteasome core complex assembly **(Figure 4C; Supplemental file S1)**, and several F-box and RING/U-box genes were differentially expressed **(Supplemental Figures S2 and S3)**. The ubiquitin–proteasome system is an important regulator of plant growth and development, and F-box and RING/U-box proteins participate in diverse regulatory processes (Sadanandom et al., 2012; Morreale and Walden, 2016; Shu and Yang, 2017; Zheng and Shabek, 2017; Ban and Estelle, 2021; Hina et al., 2024). However, the specific F-box and RING/U-box genes identified in this study have not been established as regulators of JA signaling in soybean. Moreover, the differential expression of RING/U-box genes was not uniform, with both up- and downregulated members identified. Therefore, these transcriptional changes are best considered as evidence of altered ubiquitin-related regulatory processes rather than direct evidence of altered JA signaling.

The potential relationship between JA signaling and ubiquitin-mediated proteolysis is well established through the SCF^COI1^ complex, in which the F-box protein COI1 functions acts as the receptor component required for JA-Ile-dependent degradation of JAZ repressors (Katsir et al., 2008; Santner and Estelle, 2010; Durand et al., 2016). Here, the F-box genes identified as differentially expressed in *SDP1*-suppressed seeds were not identified as *COI1* itself, and their functional relationship to JA signaling in soybean remains unknown. Similarly, changes in RING/U-box gene expression cannot be assumed to indicate altered JA signaling because these E3 ligases regulate numerous cellular and developmental pathways. Thus, although the concurrent changes in JA-related metabolites and ubiquitin-related genes are notable, additional functional studies will be required to determine whether any of these ubiquitin pathway components contribute to JA regulation in *SDP1*-suppressed seeds.

### Late-stage suppression of gluconeogenesis prevents lipid turnover

Recent studies indicate that seeds undergo beta-oxidation during development (Koley et al., 2025; Kambhampati et al., 2026) and more broadly other systems regularly recycle macromolecules as a part of cellular homeostasis in different conditions (Koley et al., 2026). Some levels of lipid turnover in seeds were suggested decades ago (Goepfert and Poirier, 2007; Graham and Eastmond 2002). Prior work in soybean suggested that gluconeogenesis may increase during late seed development, when sugar is no longer supplied by the phloem, and is tightly coordinated with lipid turnover (Kambhampati et al., 2021). Based on those observations, we proposed that fatty acids released through TAG hydrolysis could directly or indirectly help sustain carbohydrate production when a phloem source of carbon was absent (Aznar-Moreno et al., 2022; Mukherjee et al., 2025). Suppression of *SDP1* could therefore maintain the lipid sink and limit available carbon for respiration, or carbohydrate biosynthesis, including RFO production. In this study, GO-annotation of transcriptomes revealed that downregulated genes were enriched for acetyl-CoA metabolic processes and gluconeogenesis during late seed development **(Figure 4C and Supplemental file S1)**. A reduction in PEPCK activity, a crucial step in gluconeogenesis, was observed across development in the RNAi genotypes **(Figure 5)** suggesting that suppression of *SDP1* may restrict gluconeogenic carbon flux during seed development.

Collectively, our findings demonstrate that seed-specific suppression of *SDP1* enhances seed size, increases oil accumulation, and improves seed yield-related traits through changes in developmental and metabolic processes. Beyond its established role in TAG degradation, *SDP1* suppression alters carbon partitioning, cellular remodeling, phytohormone homeostasis, and multiple regulatory pathways during seed development. These integrated changes are associated with greater oil content and larger seeds without compromising seed number per plant. Therefore, seed-specific suppression of *SDP1* represents a promising biotechnological strategy for simultaneously improving seed oil content and agronomic performance in soybean.

## Author Contributions

The experimental plan was conceived by TM, BM, TPD, and DKA. Phenotyping experiments were performed by TM, SC, KG and KMM. AK and KC performed microscopy imaging. Analysis of microscopic data was performed by TM and KG. TM, KLC, ML, RW, AD performed biomass, enzyme and hormone analysis with mass spectrometry. AD, BM, and TM performed RNAseq analysis. The initial draft was composed by TM and DKA with subsequent help from TPD and BM and additional comments provided, and changes incorporated from all others. All authors approved the submission of the final version of the manuscript.

## Data Availability

All data presented in this manuscript is provided through the manuscript and supplements. Raw files for mass spectrometry and microscopy have been deposited in Zenodo (10.5281/zenodo.23378262). Raw RNAseq FASTQ files are uploaded to NCBI SRA under PRJNA1556110.

## Acknowledgements

We wish to thank the Phenotyping Facility (RRID:SCR_019049) and the Advanced Bioimaging Facility (RRID:SCR_018951) at the Donald Danforth Plant Science Center for support with imaging and image analysis of seeds, the Bioanalytical Chemistry Facility (RRID:SCR_001047) that helped with hormone analysis and provided mass spectrometry instrumentation for metabolite analyses, and the Plant Growth Facility (RRID:SCR_024902) where soybeans were grown for all experiments. We also like to thank Ms. Jiahong Zhou and Ms. Courtney Harmon for her valuable insight related to correlation analysis and cell counting. Funding was provided by the United Soybean Board, grant no. 24-203-S-B-1-A, 26-203-S-B-1-A and 27-203-A-1-B. In addition, funding from the National Science Foundation (DBI-2242822), the United States Department of Agriculture National Institute of Food and Agriculture grant no. (2023-67017-39419) and the Department of Energy Office of Science, Office of Biological and Environmental Research (DE SC0023142).

## Abbreviations

GmSDP1: Glycine max Sugar-Dependent 1
RFO: Raffinose Family Oligosaccharide
TAG: Triacylglycerol
RNAi: RNA interference
WT: Wild-type

---

**Supplementary figure S1.**
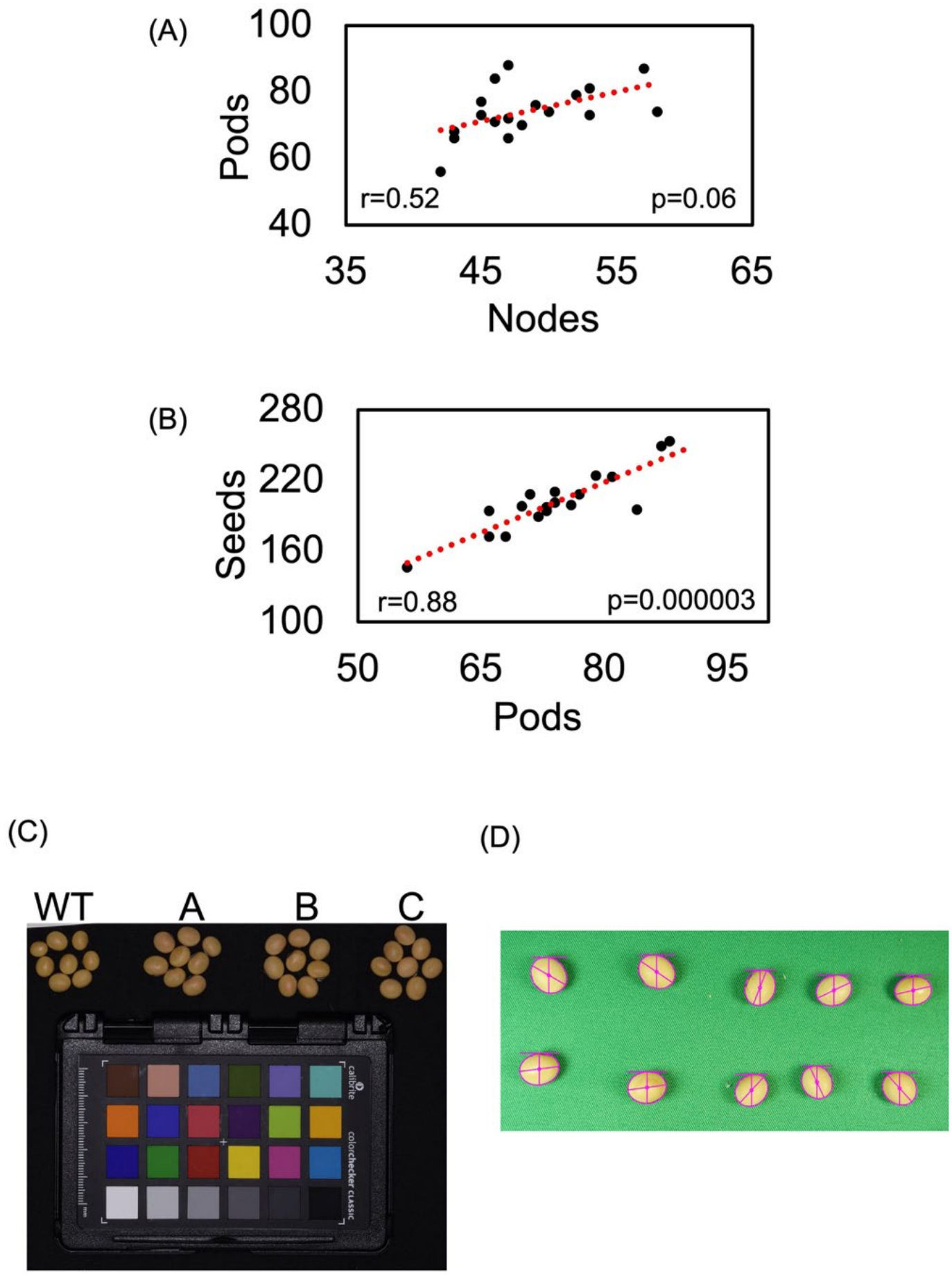
Phenotypic and agronomic measurements of WT and *SDP1-* suppressed soybean genotypes. The correlation between total number of pods and nodes (A) and total numbers of seeds and pods (B) was determined using the Pearson correlation coefficient. P-value was obtained using the corresponding t-distribution of the Pearson correlation coefficient. Mature seed images were taken using Nikon D7200 camera and a color checker card is introduced as reference for color correction and white balance (C) Seed area calculation using the morphology package in PlantCV after masking the seed from the background (D)

**Supplementary figure S2.**
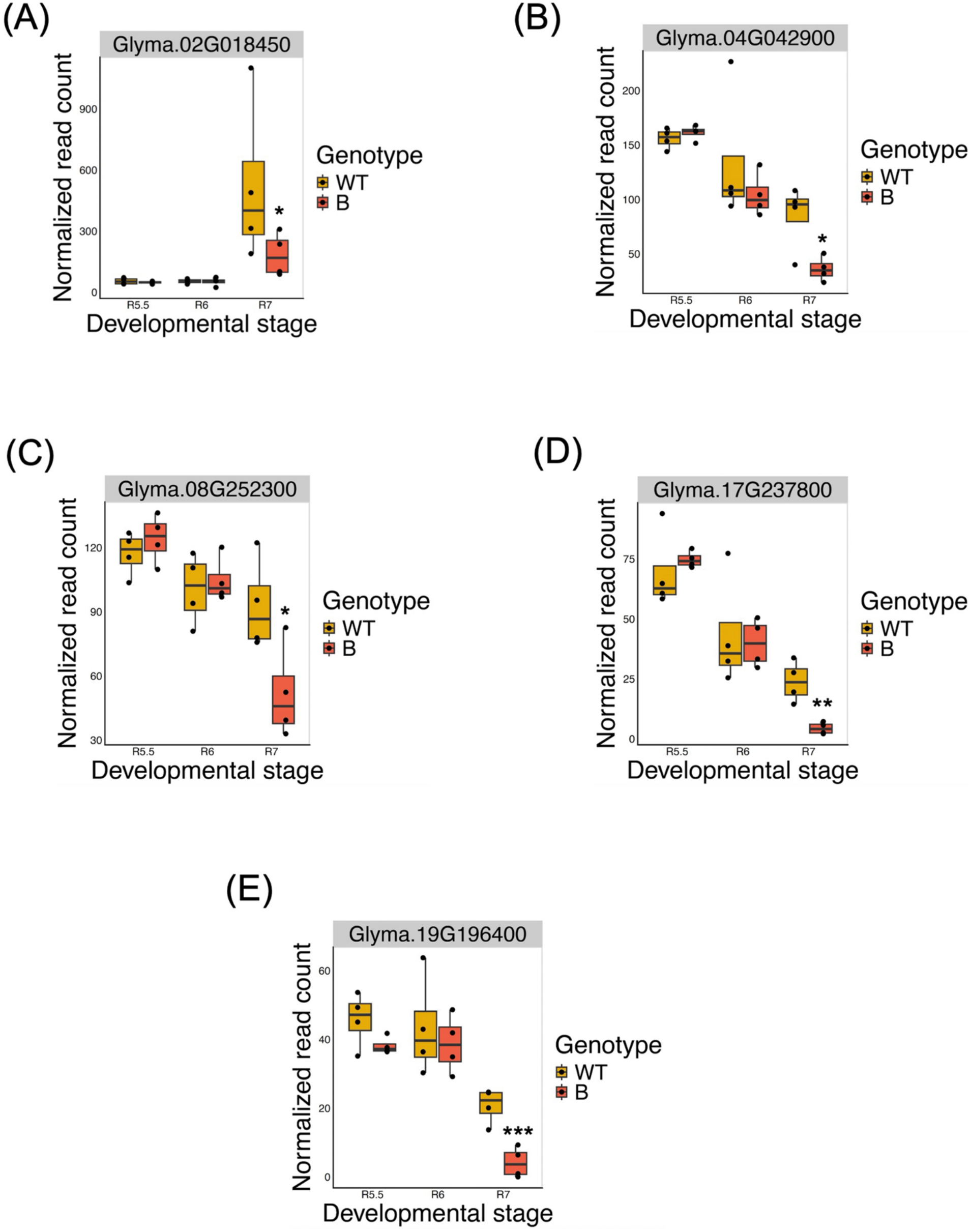
Across development expression of F-box family protein Soybean seeds from three to four plants per three reproductive stages (R5.5, R6 and R7) from WT and the transgenic genotype B (*sdp1-3, RNAi D*), was used to determine transcriptomic responses to *SDP1* suppression. Expression of five family members of the F-box family protein Glyma.02G018450 (A), Glyma.04G042900 (B), Glyma.08G252300 (C), Glyma.17G237800 (D) and Glyma.19G196400 (E). Each point represents one sample in the RNA-seq dataset; boxplots show the interquartile range, and whiskers extend to the most extreme non-outlier values. Log2 fold changes were shrunk using the adaptive shrinkage estimator (ashr; Stephens, 2016) implemented in lfcShrink(). *p∼adj∼ ≤ 0.05, **p∼adj∼ ≤ 0.01, ***p∼adj∼ ≤ 0.001, based on DESeq2 Wald test p-values adjusted for multiple testing (Benjamini-Hochberg).

**Supplementary figure S3.**
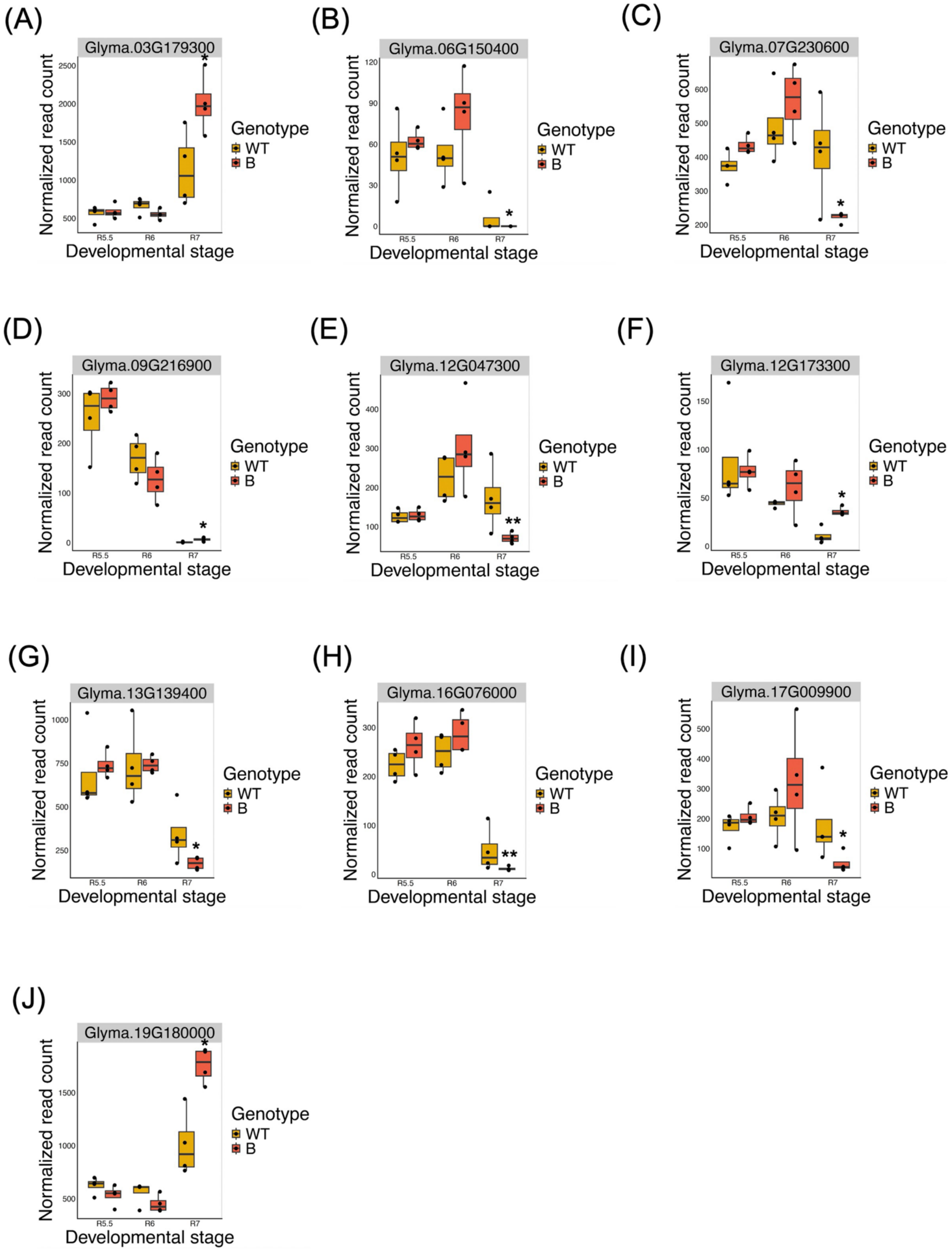
Across development expression of RING/U-box superfamily protein Soybean seeds from three to four plants per three reproductive stages (R5.5, R6 and R7) from WT and the transgenic genotype B (*sdp1-3, RNAi D*), was used to determine transcriptomic responses to *SDP1* suppression. Expression of ten family members of the RING/U-box superfamily protein Glyma.03G179300 (A), Glyma.06G150400 (B), Glyma.07G230600 (C), Glyma.09G216900 (D), Glyma.12G047300 (E), Glyma.12G173300 (F), Glyma.13G139400 (G), Glyma.16G076000 (H), Glyma.17G009900 (I) and Glyma.19G1800000 (J). Each point represents one sample in the RNA-seq dataset; boxplots show the interquartile range, and whiskers extend to the most extreme non-outlier values. Log2 fold changes were shrunk using the adaptive shrinkage estimator (ashr; Stephens, 2016) implemented in lfcShrink(). *p∼adj∼ ≤ 0.05, **p∼adj∼ ≤ 0.01, based on DESeq2 Wald test p-values adjusted for multiple testing (Benjamini-Hochberg).

**Supplementary figure S4.**
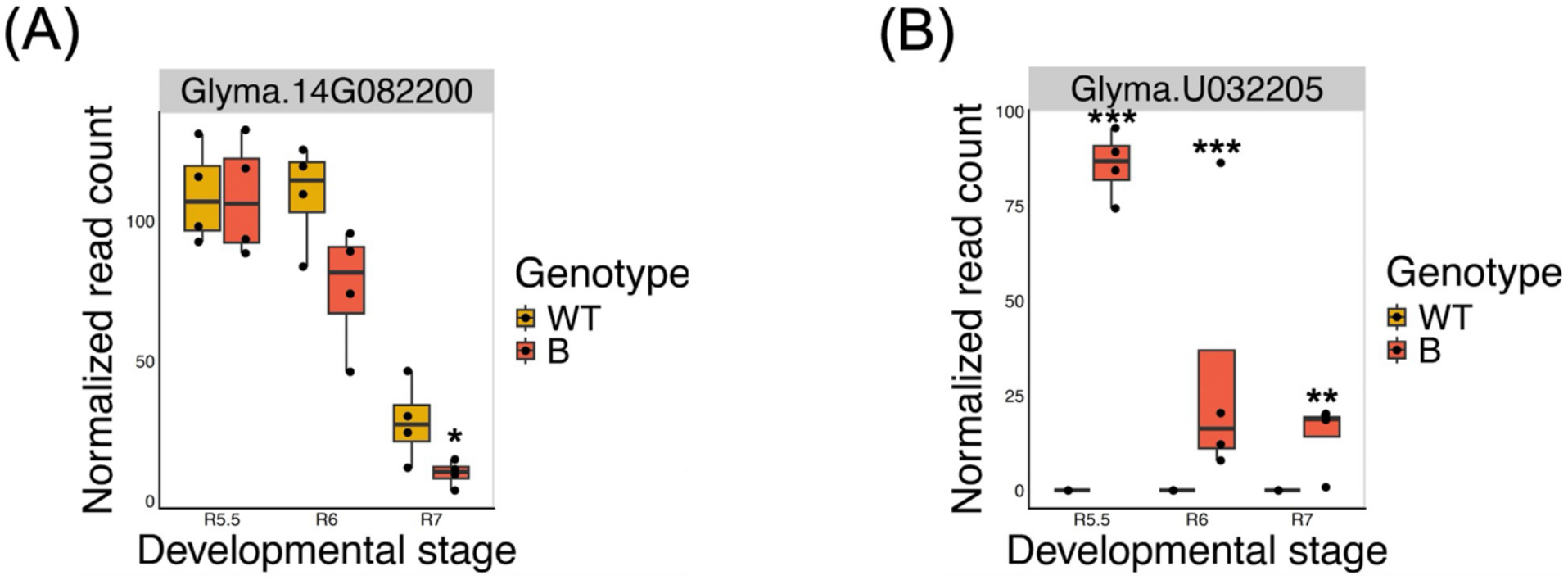
Across development expression of two Protein phosphatase 2C family protein Soybean seeds from three to four plants per three reproductive stages (R5.5, R6 and R7) from WT and the transgenic genotype B (*sdp1-3, RNAi D*), was used to determine transcriptomic responses to *SDP1* suppression. Expression of two family members of the Protein phosphatase 2C family protein Glyma.14G082200 (A) and Glyma.U032205 (B). Each point represents one sample in the RNA-seq dataset; boxplots show the interquartile range, and whiskers extend to the most extreme non-outlier values. Log2 fold changes were shrunk using the adaptive shrinkage estimator (ashr; Stephens, 2016) implemented in lfcShrink(). *p∼adj∼ ≤ 0.05, ***p∼adj∼ ≤ 0.001, based on DESeq2 Wald test p-values adjusted for multiple testing (Benjamini-Hochberg).

<u>Supplementary table S1</u>

Biomass composition of *SDP1-* suppressed soybean seeds

Soybean seed storage biomass was quantified from three to four plants per three reproductive stages and maturity (n=3-4 with 2-4 technical replicates) from WT and three transgenic genotypes A (*sdp1-1 RNAi A*), B (*sdp1-3, RNAi D*) and C (*sdp1-3, RNAi C*). Percent Biomass lipid, starch, and oligosaccharides (raffinose and stachyose) were determined. Data represents mean ± Standard error. *p≤0.05, **p≤0.01, and ***p≤0.001, indicate statistically significant difference based on a two-tailed Student’s t-test.

<u>Supplementary table S2</u>

Fatty acid composition of WT and *SDP1-*suppressed seeds

Fatty acid composition—including palmitic acid (C16:0), stearic acid (C18:0), oleic acid (C18:1), linoleic acid (C18:2), and linolenic acid (C18:3)—was analyzed from three to four plants at each of three reproductive stages plus maturity (n = 3–4, with 2–4 technical replicates each) for WT and three transgenic genotypes: A (*sdp1-1 RNAi A*), B (*sdp1-3, RNAi D*) and C (*sdp1-3, RNAi C*). Results are expressed as mol %. Data represent mean ± standard error. *p ≤ 0.05, **p ≤ 0.01, and **p ≤ 0.001 indicate statistically significant differences based on a two-tailed Student’s t-test.

<u>Supplementary table S3</u>

Percent storage reserve content change (mg/seed basis) in the seeds of three *SDP1-* suppressed genotypes: A (*sdp1-1 RNAi A*), B (*sdp1-3, RNAi D*) and C (*sdp1-3, RNAi C*) as compared to WT at maturity (R8) indicating the overall oil yield gain due to enhanced seed size.

<u>Supplemental file_S1</u>

GO annotation and DEGs of transcriptomes across developing soybean seeds from genotype B versus WT

<u>Supplemental file_S2</u>

Absolute quantification of hormone (ng/g sample) of R6 and R7 seeds from three *SDP1-* suppressed genotypes: A (*sdp1-1 RNAi A*), B (*sdp1-3, RNAi D*) and C (*sdp1-3, RNAi C*) as compared to WT

<u>Supplemental file_S3</u>

Percent lignocellulose content change in the seeds of three *SDP1-* suppressed genotypes: A (*sdp1-1 RNAi A*), B (*sdp1-3, RNAi D*) and C (*sdp1-3, RNAi C*) as compared to WT at maturity (R8)

