## Supplemental Materials and Methods for "Large seed size accompanying *GmSDP1* suppression is linked to cellular remodeling and phytohormone regulation"

**Supplemental Material and Methods:**

**Table 1**

Selected Reaction Monitoring (SRM) parameters for oligosaccharides and the ribitol internal standard on TSQ Altis mass spectrometer

| Compound | Retention ion time (min) | RT window (min) | Precursor  (m/z) | Product  (m/z) | Collision Energy  (V) | Min dwell time (ms) | RF Lens (V) |
| --- | --- | --- | --- | --- | --- | --- | --- |
| Ribitol | 7 | 14 | 151 | 89 | 18 | 131.264 | 47 |
| Raffinose | 7 | 14 | 503 | 221 | 46 | 131.46 | 141 |
| Stachyose | 7 | 14 | 665 | 383 | 50 | 131.264 | 161 |

*Phytohormone extraction*

*Materials*

Analytical reference standards for phytohormone quantification included indole-3-acetic acid (IAA; Sigma-Aldrich, St. Louis, MO, USA), N-(3-indolylacetyl)-DL-aspartic acid (IAA-Asp; Sigma-Aldrich, St. Louis, MO, USA), (±)-jasmonic acid (JA; Tokyo Chemical Industry Company, Tokyo, Japan), salicylic acid (SA; Acros Organics, Geel, Belgium), (±)-abscisic acid (ABA; Sigma-Aldrich, St. Louis, MO, USA), N-jasmonyl-L-isoleucine (JA-Ile; Toronto Research Chemicals, Toronto, ON, Canada), 12-oxo-phytodienoic acid (OPDA; Cayman Chemical, Kalamazoo, MI, USA), cis-zeatin (cZ; Santa Cruz Biotechnology, Santa Cruz, CA, USA), trans-zeatin (tZ; Caisson Labs, Smithfield, UT, USA), DL-dihydrozeatin (DHZ; Research Products International, Mount Prospect, IL, USA), and trans-zeatin riboside (tZR; Gold Biotechnology, St. Louis, MO, USA). Stable isotope-labeled internal standards included d5-JA (Tokyo Chemical Industry Company, Tokyo, Japan), d5-IAA (CDN Isotopes, Pointe-Claire, QC, Canada), d5-dinor-OPDA (Cayman Chemical, Kalamazoo, MI, USA), d4-SA (CDN Isotopes, Pointe-Claire, QC, Canada), d6-ABA (ICON Isotopes, Dexter, MI, USA), d5-trans-zeatin and d5-trans-zeatin riboside (OlChemim.cz, Olomouc, Czech Republic), ¹³C₆¹⁵N-JA-Ile, and ¹³C₄¹⁵N-IAA-Asp (New England Peptide, Gardner, MA, USA). LC–MS grade methanol (MeOH) and acetonitrile (ACN) were obtained from J.T. Baker (Avantor Performance Materials, Radnor, PA, USA), and LC–MS grade water was purchased from Honeywell Research Chemicals (Mexico City, Mexico). Standard and internal standard stock solutions were prepared in 50% (v/v) methanol and stored at −80 °C. Calibration standard solutions were prepared fresh in 30% (v/v) methanol prior to analysis.

**Table 2**

Selected Reaction Monitoring (SRM) parameters for hormones on TSQ Altis mass spectrometer

| Compound | Retention ion time (min) | RT window (min) | Precursor  (m/z) | Product  (m/z) | Collision Energy  (V) | Min dwell time (ms) | RF Lens (V) |
| --- | --- | --- | --- | --- | --- | --- | --- |
| c/t-z | 5.25 | 5.5 | 220.2 | 119.1 | 55 | 31.749 | 52 |
|  | 5.25 | 5.5 | 220.2 | 136.1 | 30 | 31.749 | 52 |
| DHZ | 5.25 | 5.5 | 222.2 | 119.1 | 55 | 31.749 | 57 |
|  | 5.25 | 5.5 | 222.2 | 136.1 | 30 | 31.749 | 57 |
| d5-tZ | 5.25 | 5.5 | 225.2 | 119.1 | 55 | 31.749 | 52 |
|  | 5.25 | 5.5 | 225.2 | 137.1 | 30 | 31.749 | 52 |
| ZR | 5.25 | 5.5 | 352.3 | 136.1 | 51.6 | 31.749 | 67 |
|  | 5.25 | 5.5 | 352.3 | 220.1 | 29.1 | 31.749 | 67 |
| d5-tZR | 5.25 | 5.5 | 357.3 | 137.1 | 55 | 31.749 | 67 |
|  | 5.25 | 5.5 | 357.3 | 225.1 | 30.4 | 31.749 | 67 |
| IAA | 11 | 6 | 176.1 | 103.1 | 49.8 | 16.586 | 41 |
|  | 11 | 6 | 176.1 | 130.1 | 25.1 | 16.586 | 41 |
| d5-IAA | 11 | 6 | 181.1 | 106.1 | 48.7 | 16.586 | 41 |
|  | 11 | 6 | 181.1 | 134.1 | 26.4 | 16.586 | 41 |
| IAA-Asp | 11 | 6 | 291.3 | 130.1 | 30.7 | 16.586 | 45 |
|  | 11 | 6 | 291.3 | 134.1 | 18.4 | 16.586 | 45 |
| ^13^C^15^N-IAA-Asp | 11 | 6 | 296.1 | 130.1 | 36.8 | 16.586 | 45 |
|  | 11 | 6 | 296.1 | 139.1 | 19.3 | 16.586 | 45 |
| SA | 11 | 6 | 137 | 65.1 | 41.6 | 16.586 | 45 |
|  | 11 | 6 | 137 | 93 | 23.7 | 16.586 | 45 |
| d4-SA | 11 | 6 | 141.1 | 69 | 47.1 | 16.586 | 45 |
|  | 11 | 6 | 141.1 | 97.1 | 25.4 | 16.586 | 45 |
| JA | 13 | 6 | 209.1 | 41 | 60 | 16.586 | 41 |
|  | 13 | 6 | 209.1 | 59 | 20.2 | 16.586 | 41 |
| d5-JA | 13 | 6 | 214.1 | 42.1 | 60 | 16.586 | 41 |
|  | 13 | 6 | 214.1 | 62.1 | 21.3 | 16.586 | 41 |
| ABA | 13 | 6 | 263.1 | 153.1 | 16.5 | 16.586 | 43 |
|  | 13 | 6 | 263.1 | 219.1 | 19.5 | 16.586 | 43 |
| d6-ABA | 13 | 6 | 269.2 | 159.1 | 17.2 | 16.586 | 43 |
|  | 13 | 6 | 269.2 | 225.2 | 21.2 | 16.586 | 43 |
| JA-Ile | 15 | 6 | 322.2 | 130.1 | 32.7 | 16.586 | 67 |
|  | 15 | 6 | 322.2 | 172.1 | 23.6 | 16.586 | 67 |
| ^13^C^15^N-JA-Ile | 15 | 6 | 329.3 | 137.1 | 32 | 16.586 | 67 |
| d5-dinor-OPDA | 22.5 | 15 | 268.2 | 170.1 | 22.8 | 26.973 | 73 |
|  | 22.5 | 15 | 268.2 | 196.1 | 28 | 26.973 | 73 |
| OPDA | 22.5 | 15 | 291.2 | 165.2 | 28.6 | 26.973 | 73 |
|  | 22.5 | 15 | 291.2 | 247.3 | 26.1 | 26.973 | 73 |
