## Supplemental Tables for "Large seed size accompanying *GmSDP1* suppression is linked to cellular remodeling and phytohormone regulation"

**Supplementary table S1**

Biomass composition of *SDP1*-suppressed soybean seeds

Soybean seed storage biomass was quantified from three to four plants per three reproductive stages and maturity (n=3-4 with 2-4 technical replicates) from WT and three transgenic genotypes A (*sdp1-1 RNAi A*), B (*sdp1-3, RNAi D*) and C (*sdp1-3, RNAi C*). Percent Biomass lipid, starch, and oligosaccharides (raffinose and stachyose) were determined. Data represents mean ± Standard error. *p≤0.05, **p≤0.01, and ***p≤0.001, indicate statistically significant difference based on a two-tailed Student’s t-test.

| Developmental Stages | Genotype | Storage reserves (% Biomass) | | | |
| --- | --- | --- | --- | --- | --- |
|  |  | Lipid | Starch | Raffinose | Stachyose |
| R5.5 | WT | 15.27±0.83 | 14.42±0.43 | 0.00±0.00 | 0.00±0.00 |
|  | A | 17.22±0.21 ******* | 13.29±0.34 ****** | 0.00±0.00 | 0.00±0.00 |
|  | B | 15.35±1.00 | 13.40±0.43 ***** | 0.00±0.00 | 0.00±0.00 |
|  | C | 15.44±0.43 | 12.52±0.28 ******* | 0.00±0.00 | 0.00±0.00 |
| R6 | WT | 24.40±0.46 | 10.84±0.42 | 0.04±0.01 | 0.02±0.02 |
|  | A | 24.02±0.55 | 10.64±0.44 | 0.04±0.02 | 0.01±0.01 |
|  | B | 26.03±0.28 ****** | 10.58±0.64 | 0.07±0.02 | 0.03±0.03 |
|  | C | 24.19±0.77 | 10.65±0.29 | 0.07±0.02 | 0.02±0.02 |
| R7 | WT | 23.97±0.33 | 2.89±0.22 | 0.70±0.03 | 3.08±0.25 |
|  | A | 24.49±1.76 | 2.35±0.45 | 0.66±0.05 | 3.51±0.41 |
|  | B | 26.91±1.01 ******* | 1.41±0.11 ******* | 0.91±0.05 ******* | 4.15±0.15 ******* |
|  | C | 26.44±0.48 ******* | 2.06±0.38 ****** | 0.93±0.13 ****** | 3.85±0.42 ***** |
| R8 | WT | 21.44±0.16 | 0.28±0.03 | 0.81±0.05 | 3.57±0.12 |
|  | A | 22.87±0.29 ******* | 0.24±0.01 | 0.75±0.04 | 3.25±0.18 ***** |
|  | B | 22.88±0.73 ****** | 0.21±0.03 ***** | 0.76±0.01 | 2.24±0.10 ******* |
|  | C | 23.04±0.14 ******* | 0.24±0.02 | 0.70±0.02 ****** | 1.97±0.06 ******* |

**Supplementary table S2**

Fatty acid composition of WT and *SDP1-* suppressed seeds

Fatty acid composition—including palmitic acid (C16:0), stearic acid (C18:0), oleic acid (C18:1), linoleic acid (C18:2), and linolenic acid (C18:3)—was analyzed from three to four plants at each of three reproductive stages plus maturity (n = 3–4, with 2–4 technical replicates each) for WT and three transgenic genotypes: A (*sdp1-1 RNAi A*), B (*sdp1-3, RNAi D*) and C (*sdp1-3, RNAi C*). Results are expressed as mol %. Data represent mean ± standard error. *p ≤ 0.05, **p ≤ 0.01, and **p ≤ 0.001 indicate statistically significant differences based on a two-tailed Student’s t-test.

| Developmental Stages | Genotype | Fatty acid (mol%) | | | | |
| --- | --- | --- | --- | --- | --- | --- |
|  |  | C16:0 | C18:0 | C18:1 | C18:2 | C18:3 |
| R5.5 | WT | 17.89±0.39 | 4.62±0.07 | 18.09±0.42 | 43.64±0.61 | 15.76±0.57 |
|  | A | 16.59±0.16 *** | 4.59±0.05 | 20.49±0.38 *** | 44.42±0.38 | 13.91±0.22 *** |
|  | B | 18.41±0.52 | 4.88±0.16 ** | 18.24±1.17 | 43.25±1.23 | 15.23±0.93 |
|  | C | 19.69±0.54 *** | 5.53±0.15 *** | 18.82±0.64 | 40.67±1.15 *** | 15.29±1.11 |
| R6 | WT | 13.68±0.32 | 4.60±0.20 | 19.73±0.19 | 52.61±0.73 | 9.38±0.14 |
|  | A | 13.44±0.20 | 4.42±0.09 | 20.74±0.51 *** | 52.29±0.51 | 9.11±0.20 |
|  | B | 12.84±0.10 ** | 4.25±0.16 | 19.46±0.75 | 55.08±0.35 *** | 8.37±0.27 *** |
|  | C | 13.27±0.17 | 4.79±0.28 | 20.37±1.07 | 52.97±1.01 | 8.60±0.31 *** |
| R7 | WT | 13.46±0.22 | 4.41±0.23 | 19.84±1.57 | 54.04±0.94 | 8.25±0.23 |
|  | A | 14.32±0.95 | 4.42±0.44 | 22.51±1.16 * | 51.22±1.77 * | 7.54±0.36 * |
|  | B | 12.90±0.39 | 4.09±0.19 | 18.88±0.92 | 56.13±1.26 * | 7.99±0.12 |
|  | C | 12.77±0.55 ** | 3.97±0.19 ** | 19.82±0.94 | 54.93±0.71 | 8.51±0.26 |
| R8 | WT | 13.46±0.66 | 4.36±0.30 | 20.80±1.11 | 53.41±0.60 | 7.96±0.18 |
|  | A | 12.87±0.09 | 3.76±0.05 ** | 21.14±1.13 ** | 54.12±0.91* | 8.11±0.27 |
|  | B | 12.85±0.11 | 3.86±0.12 ** | 19.44±0.28 | 55.72±0.10 *** | 8.12±0.03 |
|  | C | 13.17±0.14 | 4.19±0.12 | 19.97±0.61 | 54.58±0.51 * | 8.09±0.09 |

**Supplementary table S3**

Percent storage reserve content change (mg/seed basis) in the seeds of three *SDP1-* suppressed genotypes: A (*sdp1-1 RNAi A*), B (*sdp1-3, RNAi D*) and C (*sdp1-3, RNAi C*) as compared to WT at maturity (R8) indicating the overall oil yield gain due to enhanced seed size.

|  | Lipid | Starch | Raffinose | Stachyose |
| --- | --- | --- | --- | --- |
| R8 |  |  |  |  |
| A | 33.61 | 7.34 | 15.44 | 13.89 |
| B | 30.03 | -9.31 | 13.88 | -23.49 |
| C | 26.73 | -0.69 | 1.20 | -35.11 |
